# Whisker stimulation reinforces a resting-state network in the barrel cortex: nested oscillations and avalanches

**DOI:** 10.1101/2025.06.06.658305

**Authors:** Benedetta Mariani, Ramón Guevara, Mattia Tambaro, Marta Maschietto, Alessandro Leparulo, Stefano Vassanelli, Samir Suweis

## Abstract

The cerebral cortex operates in a state of restless activity, even in the absence of external stimuli. Collective neuronal activities, such as neural avalanches and synchronized oscillations, are also found under rest conditions, and these features have been suggested to support sensory processing, brain readiness for rapid responses, and computational efficiency. The rat barrel cortex and thalamus circuit, with its somatotopic organization for processing sensory inputs from the whiskers, provides a powerful system to explore such interplay. To characterize these resting state circuits, we perform simultaneous multi-electrode recordings in rats’ barrel cortex and thalamus. During spontaneous activity, oscillations with frequencies centered around 11 Hz are detected concomitantly with slow oscillations below 4 Hz, as well as power-law distributed avalanches. The phase of the lower-frequency oscillation appears to modulate the higher-frequency amplitude, and it has a role in gating avalanche occurrences. We then record neural activity during controlled whisker movements and observe that the 11 Hz barrel circuit active at rest is indeed the one involved in response to whisker stimulation. We finally show how a thalamic-driven firing-rate model can describe the entire phenomenology observed at resting state and predict the response of the barrel cortex to controlled whisker movement, suggesting that the same intrinsic dynamics underlying resting-state activity also shape sensory responses.

**Author Summary:** The brain is active even in the absence of external stimuli, generating complex patterns of activity that are thought to prepare it for processing incoming information. Two prominent features of this spontaneous activity are rhythmic oscillations and neuronal avalanches—bursts of activity that span a wide range of sizes and durations. However, how these patterns relate to actual sensory processing remains unclear. In this study, we investigated the rat barrel cortex, a well-characterized system for processing tactile information from whiskers. By combining electrophysiological recordings and computational modeling, we found that specific transient oscillations centered around 11 Hz are present during rest and become significantly stronger when the whiskers are stimulated. At the same time, spontaneous activity displays a rich dynamical regime with neuronal avalanches, whose timing is modulated by slower brain rhythms. Importantly, we show that a simple thalamus-driven computational model can reproduce these observations. We thus provide a minimal yet powerful model that suggests that the same rich, intrinsic dynamics underlying resting-state activity also shape sensory responses. Our results support the idea that spontaneous brain activity is not idle, but instead reflects an organized dynamical regime that facilitates efficient processing of sensory inputs.

## Introduction

The cerebral cortex exhibits a rich repertoire of dynamical states, even in the absence of external stimuli [15, 30, 67, 71, 80]. This continuous and spontaneous activity, known as the resting state, consumes a large portion of the brain’s metabolic energy [90], so it is likely to be functional [35]. Indeed, despite the lack of immediate tasks and demands, the resting state is far from idle; instead, it engages various intrinsic circuits, which are thought to serve a range of preparatory and regulatory functions [77]. These resting-state circuits may support the readiness for sensory processing and prepare the brain for rapid responses, effectively setting the stage for context-dependent behaviors [38, 52, 62]. In the cortex, certain networks that activate during rest can also show increased activity when a relevant external input arrives, suggesting that these circuits reflect an ongoing state of preparation and prediction, potentially optimized for varied functional needs of the brain [88]. Interestingly, the diversity of cortical activity at rest is consistent with a system operating near a critical point, where a balance is struck between order and disorder [10, 11, 57] and/or excitation and inhibition [1, 4, 66], allowing various collective patterns to emerge [2, 11, 57]. Neural dynamics in the cortex is often characterized by two main emergent phenomena: neural avalanches and collective oscillations. Neural avalanches are spontaneous cascades of neural activity that propagate through the network, displaying a scale-free distribution—meaning avalanches of all sizes, from small, localized events to large, network-spanning cascades, are observed [4, 26, 31, 34, 49, 60]. Importantly, neuronal avalanches have been found across diverse cortical scales [9, 42, 68] (recent reviews on this topic are [37, 59]), and lately they have been confirmed also at the spiking level in awake mice, after an appropriate temporal coarse-graining step [27]. Neural oscillations, on the other hand, are rhythmic patterns of activity across different frequency bands (e.g., theta, alpha, gamma) [45, 76] that support the coordination and synchronization of neural populations [7, 8, 28, 40]. These oscillations provide a temporal framework that structures neural activity, organizing the timing of communication across different regions of the cortex [5, 32, 47]. Research has shown that these oscillations can modulate the occurrence and structure of neural avalanches [56], especially through cross-frequency coupling mechanisms such as theta-gamma coupling [34]. Intriguingly, neural avalanches and oscillations are often considered in contradiction, as a dichotomy [50] between a scale-free phenomenon and a scale-specific one, characterized by a precise period of oscillation. However, in this work we suggest that this view of oscillations may be somewhat oversimplified, mostly driven by purely theoretical arguments rather than by experimental observations. Indeed, real, physiological oscillations are transient and display variability in their bursts’ duration, and even in the frequency of oscillation itself [3, 22].

The somatotopic organization of the rat thalamus-barrel circuit for processing sensory inputs from the whiskers provides a powerful model to explore this interplay.

By moving the whiskers back and forth, a process called whisking, rats can perceive the shape, size, and texture of nearby objects [20, 89]. This information is encoded into neural impulses and passed through the thalamus to various areas in the sensory-motor cortex [44], most importantly to barrels, located in the primary somatosensory cortex [61, 92], which have a somatotopic correspondence with the whisker on the rat’s snout. Furthermore, studies of whisker-related processing have shown that oscillatory activity in the barrel cortex may be closely tied to whisker movements, including both voluntary whisking during exploration and spontaneous twitching during rest [43, 73].

For these reasons, the rat barrel cortex provides an ideal model for studying intrinsic resting-state dynamics and investigating whether it is functionally connected to whisker-related sensory response. Importantly for the current work, the hypothesis that circuits that are active at rest could enhance information processing and functional efficiency remains a largely theoretical proposition. Specifically, there is little experimental evidence showing that spontaneous neural activity in cortical circuits is linked to specific functional tasks, such as whisker twitching in the barrel cortex. Following a similar line of reasoning, in [33], the authors addressed the barrel cortex using an air puff on the rat-snout for whisker-stimulation and performed avalanches analysis during spontaneous activity. The authors tailored their analyses to study whether differences in associated signatures of criticality of ongoing activity were related to an increase in the dynamic range. Another possible way to tackle the role of ongoing activity is to show that the collective dynamics at rest are characterized by a rich intrinsic repertoire (e.g., avalanches and transient oscillations), which is reinforced and recruited during task-related neural processing, such as actual whisker movement. This is indeed our approach, grounded in the hypothesis that spontaneous and task-related activity display similarities [77]; see [21] for a recent review on this topic.

To this end, we analyzed local field potential (LFP) and multi-units activity (MUAs) data recorded from the barrel cortex and thalamus of anesthetized rats (see Figure 1), both during spontaneous activity and after controlled whisker stimulation. We studied the frequency content of the signal using the empirical mode decomposition method (EMD), which allows us to perform a systematic frequency decomposition of the LFPs signals [39]. We then exploited MUAs to study avalanches. We showed that neural oscillations (centered around 11 Hz) and neural avalanches are functionally coupled in the rat barrel cortex and may work together to support whisker-related processing, even in the absence of direct stimulation. To substantiate such a hypothesis, we also recorded neural activity during controlled whisker movements and showed that the frequency centered around 11 Hz is strongly reinforced in the barrel cortex as a response to whisker stimulation [17, 24, 43, 78, 86]. We finally show how a thalamic-driven firing model [64] can describe the entire phenomenology observed at resting state and predict the response of the barrel cortex in the controlled whisker movement experiments.

**Figure 1.**
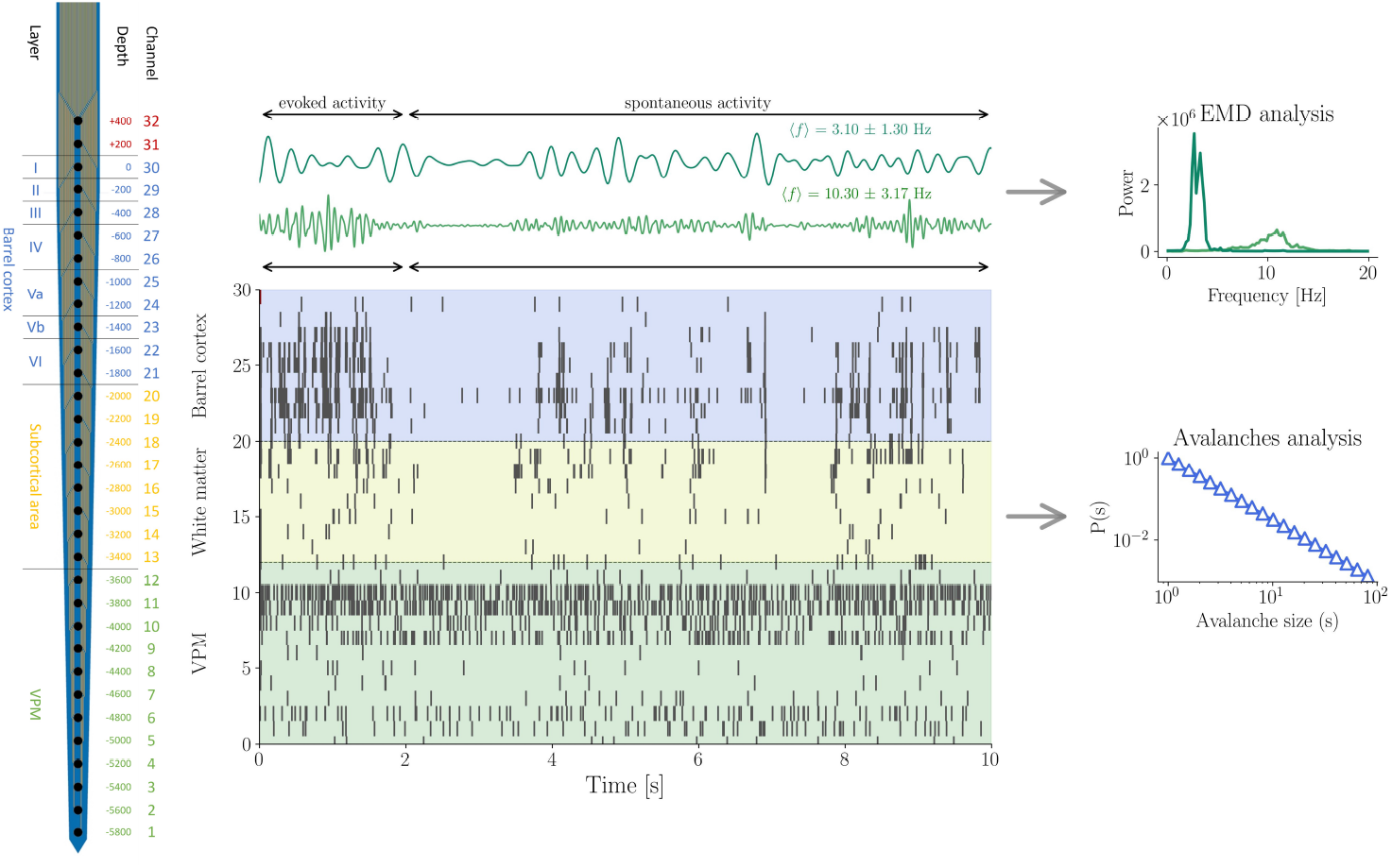
Experimental setup and outline of the analysis. **Left:** 32-channel single-shank linear probe (200 *µ*m pitch). Thirty channels were inserted into the brain, spanning the barrel cortex, subcortical regions, and part of the ventral posteromedial nucleus (VPM) of the thalamus. **Center, bottom:** Example of a single 10-s trial showing multi-unit activity (MUA) raster plots across channels (30 trials were recorded for each of the four rats). Whisker stimulation occurs at time zero. **Center, top:** The two main intrinsic mode functions (IMF 2 and IMF 4) extracted from the local field potentials are shown together with their instantaneous frequencies in the displayed trial. **Right:** Schematic representation of the analyses performed. The power of IMF 2 (fast oscillatory component) and IMF 4 (slow oscillatory component) was analyzed during both spontaneous and evoked activity in barrel cortex and thalamus channels. Avalanche analysis was performed on MUA recordings in both regions and conditions.

## 1 Results

### Resting state neural activity in the barrel cortex and thalamus

#### EMD analysis of spontaneous activity oscillations

We first studied oscillations during spontaneous (resting state) activity in the barrel cortex and thalamus (see Figure 1 for the experimental setup) in 4 rats, labeled here, in the table 1, and in the Supplementary materials, as Rat 1-4. In the current manuscript, the results presented refer to Rat 1. To characterize these oscillations during spontaneous activity we applied the empirical mode decomposition (EMD) method to the local field potentials (LFPs). This method allows for a decomposition of a signal into components that takes into account the transient and non-linear nature of physiological signals, as opposed to a decomposition into Fourier components. The components of the EMD method are called intrinsic mode functions (IMFs). We focused on the second intrinsic mode function (IMF 2) which has frequencies in the range of the whisker-stimulation response (mean ± std of the distribution = 11.13 ± 2.83 Hz) and the intrinsic mode function number 4 (IMF 4), which is a low-frequency oscillation (with frequencies mean ± std = 2.78 ± 0.70 Hz). Details on the selection of the IMFs can be found in the supplementary material, where we explain that IMF 1 is a high-frequency residual, typically excluded in the EMD analysis [13, 22]. Using the Hilbert transform [84] we obtained the Hilbert spectrum, which is a weighted non-normalized joint amplitude-frequency-time distribution [39]. In particular, here we focused on the marginal spectrum (see Methods section), obtained by integrating the Hilbert spectrum over time. We thus obtained the power of the individual IMFs (for each trial) and then extracted the values of their peaks. In Figure 2 the marginal spectrum for IMF 2 and IMF 4 in all the trials for an example rat is shown, both for the channels positioned in the barrel cortex (Fig. 2 a) and for the channels recording thalamus activity (Fig. 2 b). We made a statistical comparison (t-test) between the barrel cortex and the thalamus in terms of the intensity of the IMF power peaks. We found that power of IMF 2 peaks is significantly higher in the barrel cortex as compared to the thalamus during spontaneous activity (p-value ≪ 0.001)(see Figure 2).

**Table 1.** One-sided t-tests p-values: the peak of the marginal Hilbert spectrum of IMF 2 and IMF 4 is calculated for a barrel cortex layer IV channel (labeled as “cort.” in the table) and for a central thalamus channel (labeled as “thal.” in the table), in all the 30 trials for each rat, considering a window of 2 seconds after the stimulation as the evoked activity, and the following activity as the spontaneous activity. Paired t-test-tests are performed among the values of the spectrum peaks.

|  | Rat 1 | Rat 2 | Rat 3 | Rat 4 |
| --- | --- | --- | --- | --- |
| IMF-2 peak stim cort. vs IMF-2 peak rest cort. | pv = $2.96e^{-7}$ | $6e^{-5}$ | pv = 0.003 | pv = $7.76e^{-7}$ |
| IMF-4 peak stim cort. vs IMF-4 peak rest cort. | pv = 0.1 | pv = 0.98 | pv = 0.05 | pv = 0.24 |
| IMF-2 peak stim thal. vs IMF-2 peak rest thal. | pv = 0.37 | pv = 0.01 | pv = 0.07 | pv = 0.03 |
| IMF-4 peak stim thal. vs IMF-4 peak rest thal. | pv = 0.94 | pv = 0.74 | pv = 0.1 | pv = 0.15 |

**Figure 2.**
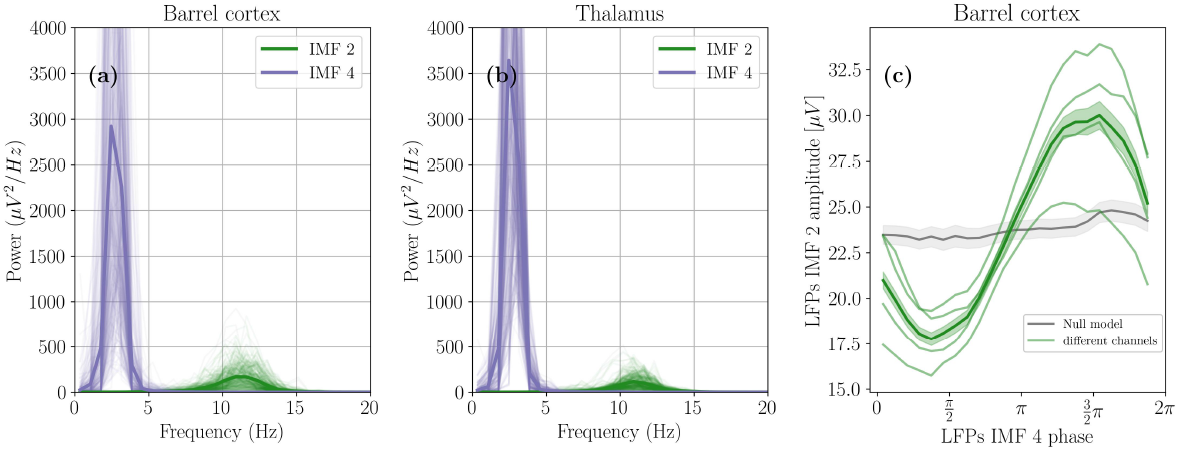
Barrel cortex and thalamus oscillatory activity during spontaneous activity. Marginal Hibert spectrum of the IMF 2 and IMF 4 rhythms during the spontaneous activity in the barrel cortex (a) and in the thalamus (b). Light-colored lines refer to the different channels either of the barrel cortex (a) or of the thalamus (b), and all the trials are also reported. The darker lines indeed refer to an average of the power across trials and channels. (c) Phase amplitude coupling between the phase of IMF 4 and the amplitude of IMF 2 of the LFP signals in the barrel cortex. Different lines represent the curve for each channel, averaged across trials. The dark line is the average curve across channels and trials, while its shaded band has a width corresponding to the standard deviation of the mean across channels and trials. As explained in the main text, the gray line is a null model, i.e., the curve in gray is obtained by randomly matching the trials of IMF 4 with the trials of IMF 2 (for each channel), and then averaging across channels and trials.

Given the presence of oscillations during resting-state periods, we explored whether the slow oscillation (IMF 4) modulates the amplitude of the faster oscillation (IMF 2) via phase-amplitude coupling (PAC). As Figure 2 (c) shows, there is a sinusoidal relationship between the phase of IMF 4 and the amplitude of IMF 2, indicative of PAC and nested oscillations [5, 47, 51] within the barrel cortex. Indeed, the presence of PAC is defined by a systematic modulation of the amplitude of the fast oscillation as a function of the phase of the slow oscillation, resulting in a non-uniform (e.g., sinusoidal) dependence, meaning that high-amplitude bursts of IMF 2 have a precise IMF 4-phase-preference. To clarify the strength of the effect, we also show the same curve for a null model (the gray line in Fig. 2c). To obtain the null model, we have matched the IMF 2 time series of each trial with the IMF 4 from a randomly selected trial, thus disrupting the temporal correspondence between the phase of the slow oscillation and the amplitude of the fast one. Note that this null model is particularly conservative, since we disrupt the two-modes-correspondence without the need to generate an artificial shuffled signal for IMF 4; instead, we consider a surrogate signal with the same autocorrelation characteristics of the original. As it is possible to see, in a scenario without modulation, the PAC curve would be a flat line corresponding to a baseline value, displaying thus no phase-preference. Remarkably, in all the animals, we find a PAC curve that is significantly far from the null model (see Supplementary Material for the results for all the animals).

#### Avalanche dynamics during spontaneous activity

We then studied the distribution of avalanches during spontaneous activity by analyzing multi-unit activity (MUA) recordings. Indeed, as opposed to LFPs, MUAs display discrete events by nature and thus allow an avalanche analysis less sensitive to signal-thresholding [53, 85, 87]. In order to detect avalanches, we employed the average inter-event interval (also called ISI, i.e., inter-spike interval) (see the Methods section for the full details). This resulted in the following ISI for avalanches analysis in the ongoing activity of the barrel cortex for the four rats: 5.37 ± 0.04 ms, 3.57 ± 0.03 ms, 4.04 ± 0.03 ms, 4.12 ± 0.03 ms; and the following ones for the thalamus: 2.74 ± 0.01 ms, 2.60 ± 0.01 ms, 1.97 ± 0.01 ms, 1.66 ± 0.004 ms. In the barrel cortex, we found power-law distributions were observed both in the duration and size of the avalanches, i.e., we found that the avalanches’ sizes were distributed as *p*(*S*) ∼ *S*^−*τ*^ and the avalanches’ durations as 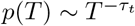 (Figure 3 (a, b)), indicative of a highly structured spatio-temporal dynamics. Interestingly, the exponent *δ*_*fit*_ linking the average size given duration ⟨*S*⟩(*T*) and the duration itself of an avalanche *T* (see inset in Figure 3b) is *δ* = 1.30 ± 0.05, close to the one observed by [26](*δ*_*fit*_ ≈ 1.28), [31] (*δ*_*fit*_ ≈ 1.3), and [53, 54] (*δ*_*fit*_ in the range 1.21-1.28). Notably, the same analysis of neuronal avalanches in the thalamus did not reveal the presence of power-law distributions. Instead, as can be seen from Figure 3 (e) and (f), the distributions are better fitted by exponentials. Also, the relation between ⟨*S*⟩ (*T*) and the duration itself of an avalanche *T* revealed a scaling that can be observed in trivial processes [54], with an exponent *δ*_*fit*_ = 1.09 ± 0.03 close to one. This means indeed that avalanche sizes are simply proportional to avalanche durations, without exhibiting any particular spatio-temporal dynamics. These results interestingly suggest that avalanches emerge in the barrel cortex and are not already present at the thalamus stage.

**Figure 3.**
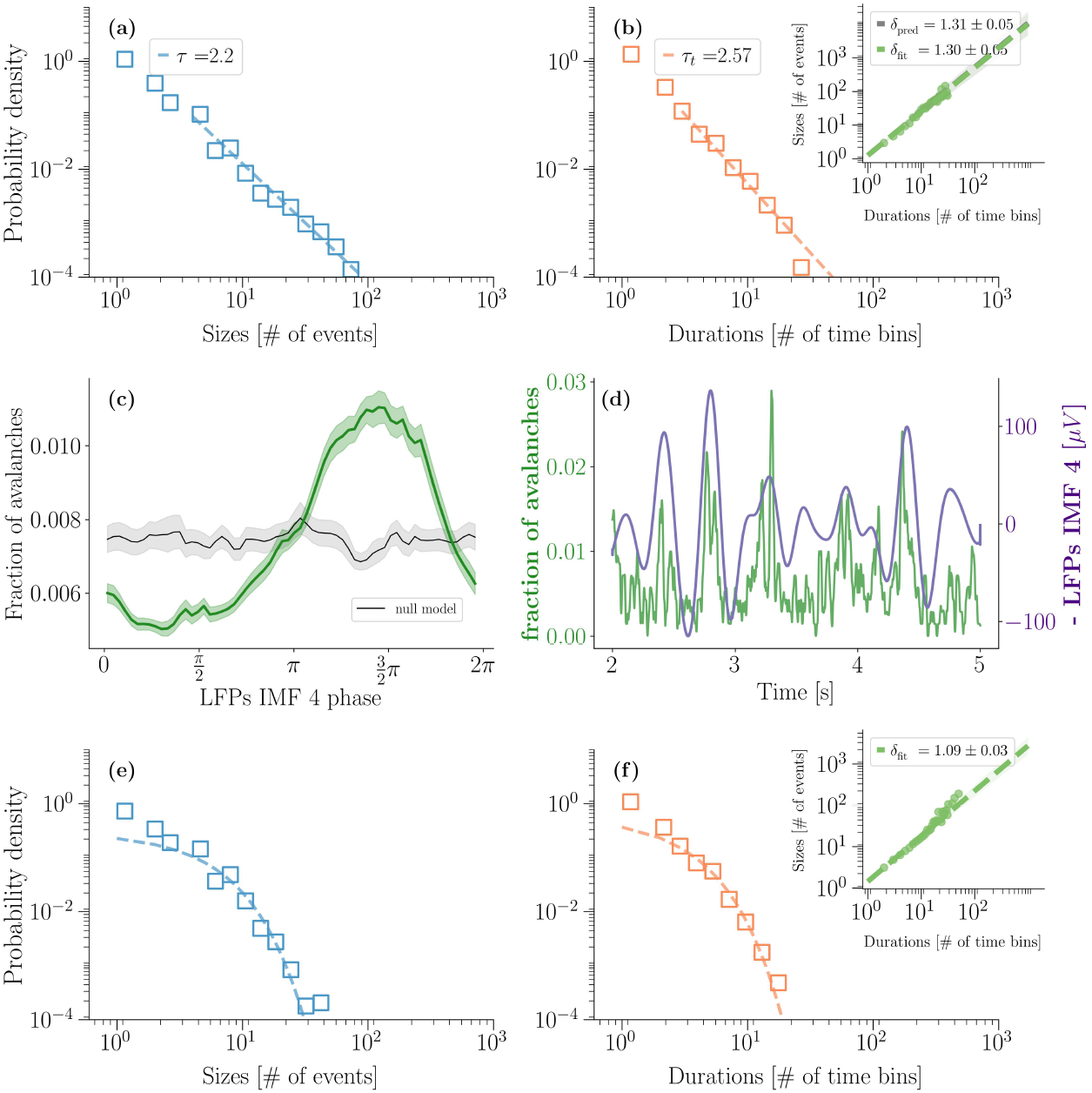
Avalanches during spontaneous activity in the barrel cortex and thalamus and coupling between the slow oscillation and the fraction of avalanches. (a, b) Avalanches’ sizes (*S*) (a) and durations (*T*) (b) distribution in the barrel cortex during spontaneous activity calculated using MUAs. In the inset of (b) it is possible to see the relation between ⟨*S*⟩ (*T*) and *T* . (c) Coupling between the phase of IMF 4, i.e. the low-frequency oscillation of the barrel signals and the fraction of avalanches in the barrel cortex. The shaded band indicates the standard deviation across trials. The black line indicates the results for a null model, which is obtained by randomly associating the trials of the fraction of avalanches’ time series with the trials of the IMF 4 phase. (d) Example of a trace of the fraction of avalanches with the respective evolution of IMF 4. Note that IMF 4 in Figure (d) has been flipped with respect to the *x* −axis since in local field potentials (LFPs), negative peaks correspond to neural population activations. In contrast, avalanche density is always expressed as a positive quantity and does not exhibit negative values. (e, f) Avalanches’ sizes (*S*) (e) and durations (*T*) (f) distribution in the thalamus during spontaneous activity calculated using MUAs. In the inset of (f) it is possible to see the relation between ⟨*S*⟩(*T*) and *T* .

In the barrel cortex, where we observe power-law distributed avalanches, we also computed avalanches’ density, defined as a measure of the presence of avalanches in a given interval of time (see Figure 3 (c, d)). We computed the coupling between the slow oscillation phase and the fraction of avalanches derived from multi-unit activity (MUA), which also follows a sinusoidal pattern consistent with the one observed in LFP signals (Figure 3 (c, d)). This suggests that avalanches in the barrel cortex consistently occur in specific phases of the slow oscillation, following the same pattern of the IMF 2 oscillation (Figure 2 (c)). Importantly, these features suggest that avalanches and fast oscillations (IMF 2) are complementary aspects of the same underlying dynamics. Indeed, the avalanche detection algorithm groups in the same avalanche oscillatory bursts (consequent oscillatory cycles) that happen across the barrel column. The slow oscillation, in turn, modulates the timing of both processes. Our results support a scenario in which scale-free avalanche dynamics coexist with, and are temporally structured by, nested oscillatory processes.

### Neural activity after whisker stimulation

#### EMD Analysis of post-stimulation oscillations

We also applied the EMD analysis in the periods following whisker stimulation, to understand its effect on IMFs 2 and 4. We again used the Hilbert-Huang transform to extract the power of the peak of each IMF considering a time interval of two seconds after the stimulation. We then performed t-tests on the IMF power peaks for statistically comparing evoked activity and ongoing activity, both in the barrel cortex and in the thalamus (Table 1 for the statistical results). Results in Table 1 indicated a significant difference in the IMF 2 band between spontaneous and post-stimulus activity within the barrel cortex. Whisker stimulation reinforces the oscillation in this frequency range, while in the thalamus, p-values are always greater than 0.01. Moreover, as found in resting state recordings, the peaks in this band were more pronounced in the barrel cortex than in the thalamus, as shown in Figure 4. In contrast, oscillations in the IMF 4 band were present consistently across both cortex and thalamus without significant differences between resting and post-stimulus states.

**Figure 4.**
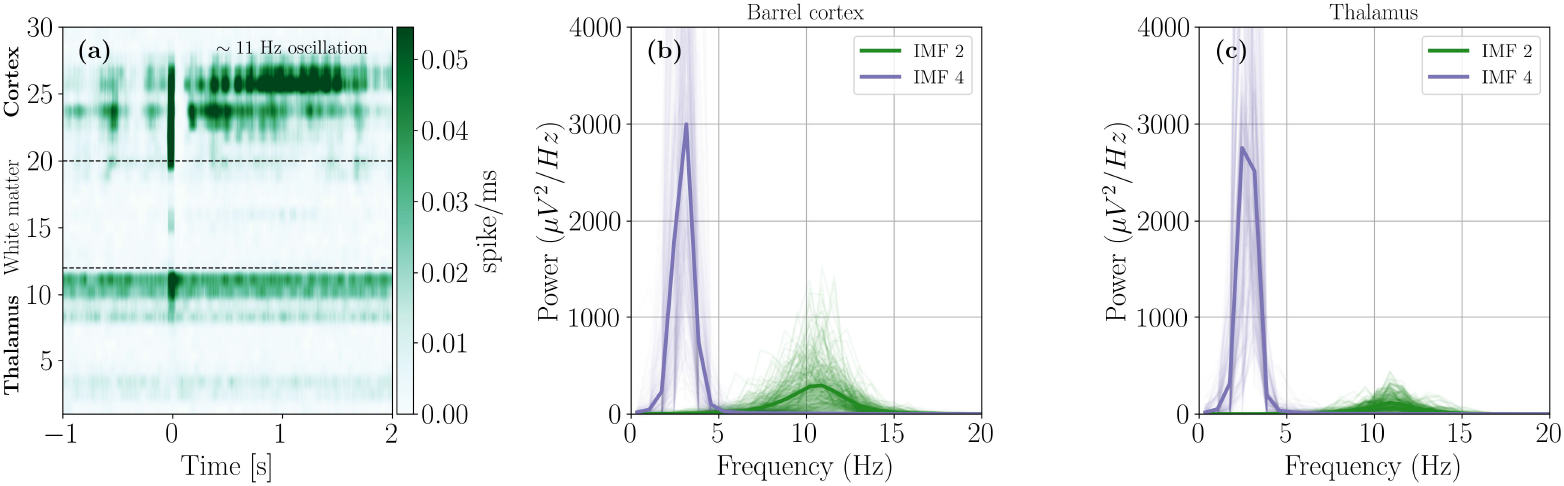
Barrel cortex and thalamus oscillatory activity after whisker stimulation. (a) Firing rate obtained from MUAs by averaging across all trials, after whisker stimulation (at time zero). Oscillatory activity at 11 Hz is observed in the barrel cortex immediately after stimulation. Such oscillation is absent in the thalamus. (b,c) Marginal Hilbert spectrum of IMF 2 and 4. As in the case of spontaneous activity, the power of the IMF 2 is significantly higher in the barrel than in the thalamus. Moreover, by comparing insets (b) and (c) with those of Figure 2, there is a significant increase in the power of IMF 2 after whisker stimulation (but see also Table 1).

#### Avalanche statistics after stimulation

After whisker stimulation, we analyzed avalanche dynamics, finding a power-law distribution in avalanche sizes in the barrel cortex, as observed during spontaneous activity (see supplementary material). However, a slight bump in the distribution was observed, corresponding to large avalanches related to the stimulus. This result is consistent with our previous finding [53]. These enhanced avalanches co-occurred with IMFs 2 oscillations in the barrel cortex (Figure 4 (b)), suggesting that post-stimulation oscillations may coexist with an increase in avalanche activity. See the Supplementary Materials for an intuition on the relationship between avalanches and the fast oscillation. On the other hand, in the thalamus, we found avalanche distributions very similar to the spontaneous activity ones, described by exponential distributions (see supplementary material).

### Modelling the neural dynamics in the barrel cortex

#### Oscillatory activity in the model in resting state

Finally, we investigated the emerging properties of the neural dynamics of the barrel cortex by using a modified version of the reduced model proposed by Pinto and Ermentrout [64, 65] schematized in Figure 5 (a). We used the same set of parameters identified experimentally in a previous study [46], which were calibrated and fitted to match the neural activity observed in the barrel cortex following whisker stimulation in rats (see Methods for details). In our work, the parameters differ only for the one that describe the cortical-thalamic connections, which are calibrated to describe our data and differ (within the experimental standard deviations provided by [64]) with those obtained in the Kyriazi and Simons study [46]. Specifically, the weight of the connection between the thalamus and the inhibitory population (*w*_*ti*_) was decreased. In such a way, while the internal circuit of the barrel cortex remains dominated by inhibition, as was the case in the original derivation of the model [64], our model contains limit cycles in its dynamical repertoire with frequencies that remarkably overlap in the range 10-15 Hz of the frequencies that we find experimentally. Interestingly, using MUAs firing rates from the thalamus at resting state as input, the model exhibits both slow oscillations (IMF 4) and faster oscillations (IMF 2), that can be observed experimentally. Figure 5 (b, e) shows the activity of the model and spectrum for IMF 2 and IMF 4 for spontaneous activity. We also computed the phase-amplitude coupling between these two modes exhibited by our model and found that they constitute nested oscillations, as observed in Figure 5 (d). Indeed, it shows a sinusoidal relationship between the phase of the slow oscillation and the amplitude of the faster oscillation, replicating our experimental results.

**Figure 5.**
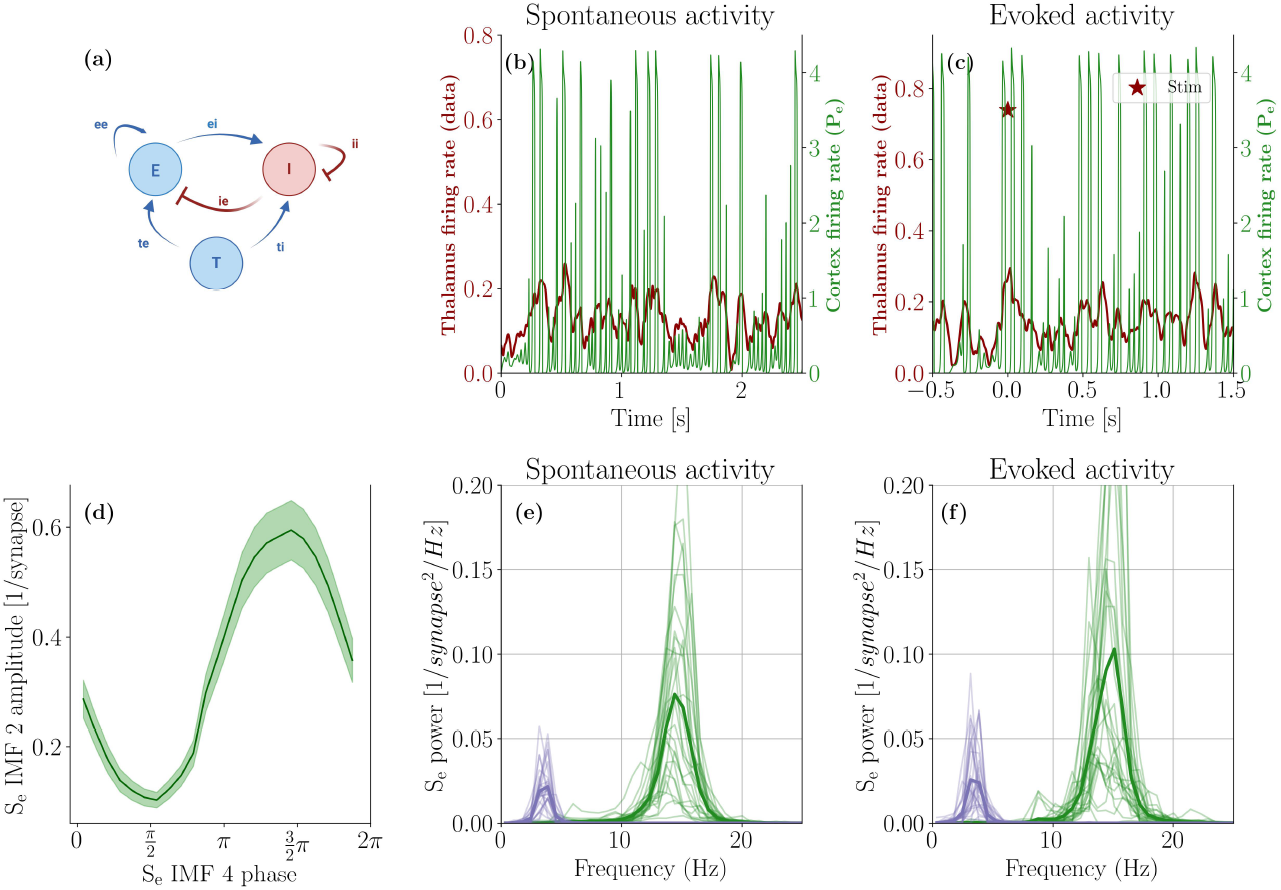
Oscillatory activity in the model. (a) Schematic illustration of the cortical barrel circuit (composed of an excitatory (E) and an inhibitory (I) subpopulation) that receives external excitatory input from the thalamus (T). (b) In dark red, the time histogram of the firing rate of the thalamus (*P*_*T*_) during spontaneous activity obtained directly from MUAs, used as input of the model. In green, the firing rate of the excitatory units of the barrel cortex (*P*_*e*_) in response to the thalamic input. (d) Phase-amplitude coupling is computed on the simulated data, which provides the relation between the phase of the low-frequency oscillation (IMF 4) and the amplitude of the high-frequency oscillation (IMF 2); note that the phase of the IMF 4 has been shifted of *π*, in order to help a comparison with Figures 2 and 3) where the IMF 4 of the Local Field Potential is considered (and thus in that case negative excursion corresponds to activation of populations of neurons). (e) Marginal Hilbert spectrum of the intrinsic mode functions (IMF) 2 and 4 of the modeled cortical excitatory population. In (c, f), the same quantities of (b, e) are shown, yet giving as input the experimental thalamus’ firing rate after a whisker stimulation.

#### Avalanche dynamics in the model

We then computed the avalanches’ also in the model, and studied coupling also as regards the fraction of avalanches. As detailed in the methods, we generated an event plot from the corresponding cortical firing rate, and then computed avalanches in the same way as they are measured experimentally. Importantly, our model is able to reproduce the barrel-cortex power-law distribution of avalanches that we observe experimentally (See Figure 6 (a) and (b)). The scaling of ⟨*S*(*T*) ⟩ as a function of *T* is also reproduced, with only very large avalanches deviating from it in the tails of the distributions [19, 53]. Then, we computed the coupling between the phase of IMF 4 of *S*^*e*^ (cortical simulated activity) and the fraction of avalanches. Our modeling framework is able to reproduce the phase preference that we observe also experimentally as regards the fraction of avalanches (see Figure 6 (c) and (d)).

**Figure 6.**
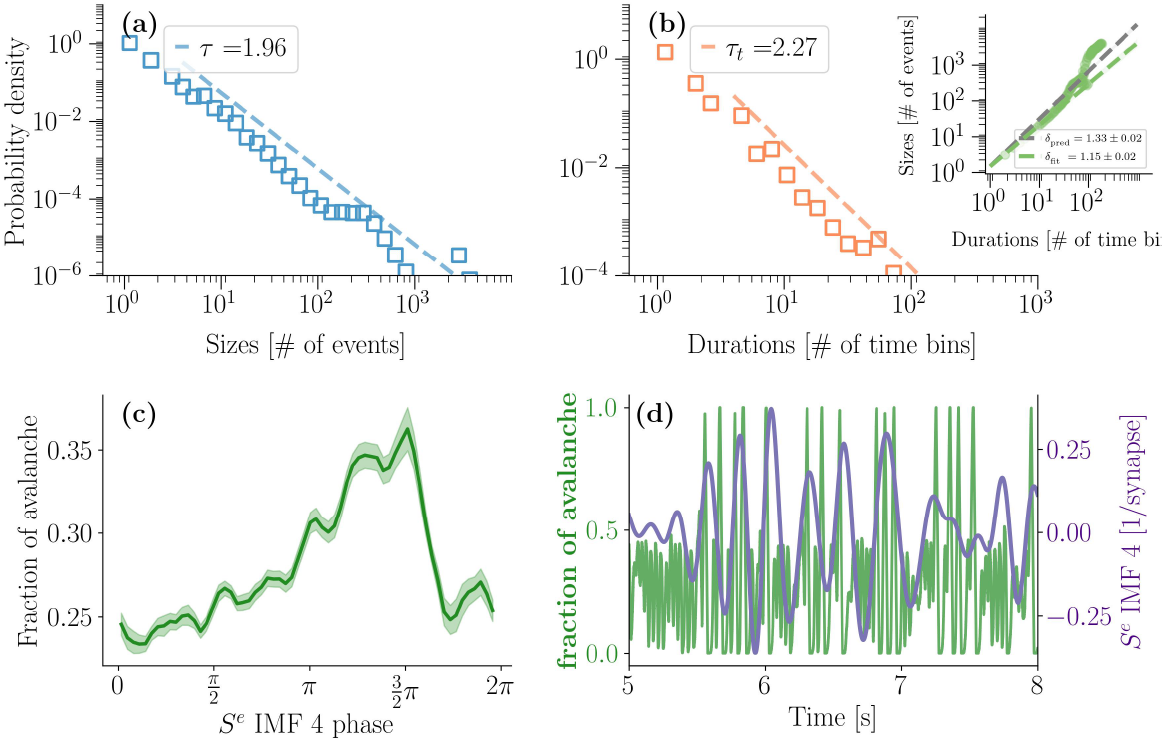
Avalanche dynamics in the model. (a, b) Distribution of avalanche sizes and durations obtained from the cortical firing rate of the model. In the inset of (b) it is possible to observe the scaling between ⟨*S*(*T*) ⟩ and *T* (mean avalanche size versus corresponding duration). (c) Coupling between the phase of IMF 4 of the synpatic drive of the cortical population, and the simulated fraction of avalanches, measured in the same way as in the experiments. (d) Example of a trace of the fraction of avalanches with the respective evolution of IMF 4.

#### Model Response to Whisker Movement

We then explored the model prediction of the neural activity after whisker movement by using as input the corresponding MUAs firing rates from the thalamus. We emphasize that we did not modify any of the parameters previously used to describe the resting state (see Table 2); the only change was the thalamic input, which we replaced with the experimentally measured firing rate following whisker stimulation (see Figure 5 (c)). Remarkably, the model predicts the enhancement of the IMF 2 oscillations (see Figure 5 (f)). We then performed a statistical (t-test) analysis of the peaks of the power of the excitatory population in the model. It indicated that the power of IMF 2 is significantly higher (p-value ≪ 0.01) in the after-stimulation period as compared to the resting state. This is in accordance with our experimental results of table 1.

**Table 2.**

| parameter | symbol | value |
| --- | --- | --- |
| resting potential (mv) | $\rho$ | -60 mV |
| firing threshold (mv) | $\theta$ | -45 mV |
| temperature | $T_e$ | 12.03 |
| - | $T_i$ | 11.62 |
| conversion factor | $\gamma_e$ | 5 |
| - | $\gamma_i$ | 15 |
| refractory parameter | $e_r$ | 2.2 |
| - | $i_r$ | 4.1 |
| connection strength (mv) | $w_{ee}$ | 1.0 |
| - | $w_{ei}$ | 1.0 |
| - | $w_{ie}$ | 2.0 |
| - | $w_{ii}$ | 1.5 |
| - | $w_{te}$ | 3.9 |
| - | $w_{ti}$ | 1.5 |
| number of synapses | $ec$ | 42 |
| - | $ic$ | 12 |
| - | $tc_e$ | 12 |
| - | $tc_i$ | 10 |
| decay constant (ms) | $\tau_e$ | 5 |
| - | $\tau_i$ | 15 |

## 2 Discussion

In the current study, we investigated oscillations and avalanches in the thalamocortical circuit of urethane-anesthetized rats, both during spontaneous activity and after mechanical stimulation of the whiskers. In both cases, a low-frequency oscillation (IMF 4, with frequencies 2.78 ± 0.70 Hz (mean ± std of the distribution)) was detected concomitantly with a high-frequency oscillation (IMF 2, 11.13 ± 2.83 Hz (mean ± std of the distribution)). When the whisker is stimulated, the power of this oscillation increases in several layers of the barrel cortex. In the thalamus, this phenomenon was not observed. In general, the power of the higher frequency oscillation was larger in the barrel cortex compared to the thalamus, both in the evoked activity and in the spontaneous activity. In contrast, the power of the low-frequency oscillation in the barrel cortex did not change after whisker stimulation. Our results show a clear phase–amplitude coupling between low- and high-frequency oscillatory activity, where the phase of the slower rhythm modulates the amplitude of faster oscillations, consistent with the concept of “nested” oscillations [5]. This is in line with prior work by Ito et al., who demonstrated modulation of gamma-band local field potential amplitudes by the respiratory rhythm [41], and with studies on the human thalamus that describe structurally constrained phase–amplitude coupling [47, 51].

We found that neuronal avalanches in the barrel cortex follow a power-law distribution, indicative of complex temporal and spatial dynamics reminiscent of criticality [19, 50]. Importantly, the phase of low-frequency oscillations modulates the density of avalanches calculated from multi-unit activity (MUA), mirroring the nesting seen in the LFPs.

By driving our model with empirical thalamic firing rates, we replicated the observed nested oscillations during rest and predicted their amplification upon stimulation. Our model is moreover able to reproduce the other resting-state phenomenologies of our experiments, i.e., the avalanches’ statistics, and the modulation of the fraction of avalanches through the slow oscillation. All these features are indicative of rich and complex spatio-temporal dynamics. Our earlier work [54] showed that critical-like dynamics, characterized by enhanced mutual information between spiking neurons across brain regions, can emerge under time-varying external inputs. In this regard, we note that our thalamo-cortical model does not operate persistently at a fixed critical point. Rather, due to the time-varying thalamic input, the cortical circuit dynamically fluctuates around the bifurcation. This feature is consistent with our experimental observations of intermittent oscillations. Indeed, our model shows that the avalanching (“critical-like”) behavior is due to the interplay of the external thalamic drive together with the intrinsic, non-linear dynamics of the barrel cortex. Both features combined lead to transient excursions above the Hopf bifurcation; these excursions end when the thalamus input decreases below the bifurcation threshold.

We hypothesize that the “critical-like” dynamics observed during resting state may provide a rich environment characterized by nested oscillations and avalanches, allowing for a flexible, yet structured, resting state network that may “prime” the cortex for rapid, coherent responses to whisker input. If this were the case, such preparatory dynamics could enhance the cortex’s ability to integrate incoming sensory information, implying that whisker-related processing is facilitated by resting-state activity. The oscillations observed after whisker stimulation were already in the intrinsic repertoire of the resting state [15, 77] and they may set a temporal scaffold within which avalanches can propagate in response to specific sensory stimuli. The interpretation we propose here is supported by our reduced excitatory-inhibitory neuronal population model.

Interestingly, often oscillations and power-law neuronal avalanches are seen in dichotomy [50], as the former are associated with a scale-specific phenomenon, described by a characteristic temporal scale, while the latter lack a characteristic scale (scale-free), as they are distinguished by power-laws. However, we believe that this contradiction is only apparent if formalized in our framework. Indeed, the oscillations displayed by our experiments, and by our model, are transient and intermittent (they do not persist for the entire recording). In other words, even though a characteristic frequency of oscillation is present (that is, however, identified through empirical mode decomposition, which allows for a flexible frequency variability in time [39]), the durations of its bursts display inherent stochasticity and variability, and, when measured in terms of avalanches, display a power-law distribution. This picture is consistent with recent (and less recent) works that indeed find no contradiction in the coexistence of oscillations, synchronizated oscillators, and power-law avalanches [6, 19, 50, 66].

Notably, spindle oscillations in the 11-15 Hz range-previously observed in barrel cortex LFPs during sleep and anesthesia [82] share this frequency band. Spindles, known to support memory consolidation [36], have also been documented during development [43] and anesthesia [18]. While we did not perform a formal spindle detection, the analyzed frequency range likely includes spindle activity.

This rhythm has been associated with resting-state synchrony, attention modulation, sensory hypersensitivity, and even seizure-like events [25, 73, 74, 78]. Spindles are believed to originate in the thalamus [58], a hypothesis consistent with our model: oscillatory cortical output emerges when thalamic input exceeds a threshold, even though the input lacks spindle-like patterns.

It is important to notice that the experiments we conducted involved anesthetized rats, and it will be of foremost importance to confirm these results also on awake animals, to avoid confounders due to the anesthesia. Moreover, given that the rats are anaesthetized, we exclude that the spontaneous activity oscillations we observe correspond to some rats’ behavior, such as, for example, spontaneous whisking. We instead believe they constitute a neural activation of the intrinsic activity repertoire available to the animals. The fact that the spindle-frequency oscillations emerge prominently after the whisker is stimulated, suggests that these oscillations are linked to a latent dynamical mode relevant for sensory inputs, rather than just an effect of anesthesia. On the other hand, we believe that there is the possibility that the oscillations with frequency below 4 Hz are affected by anesthesia [14, 79]. Indeed, urethane anesthesia is known to display UP and DOWN states [14, 26], compatible with the slow oscillations in our recordings. In the present analysis, we did not explicitly segment the recordings into distinct brain states (e.g., UP and DOWN states), and avalanche statistics were computed over the entire recording. This choice was motivated by the fact that our primary goal was to characterize the overall statistical properties of the ongoing activity, considering its temporal dynamics, rather than to isolate state-specific regimes. Also, the study of different cortical states in our case would have required a severe segmentation of the recordings, disrupting the continuity of temporal dynamics, which was our main object of interest. We acknowledge, however, that a segmentation into UP and DOWN states could reveal differences in avalanche statistics across conditions [26, 33], potentially leading to regimes that deviate from or more closely follow power-law behavior. Importantly, the analysis of oscillatory activity was specifically designed to account for transient and nonstationary dynamics, as it relies on empirical mode decomposition, which does not assume stationarity and does not average over time in the same way as classical spectral methods. This makes it well-suited to capture intermittent oscillations across varying brain states. Regarding avalanche statistics, we find that considering the full recording yields power-law-like distributions.

In [33], the authors performed an interesting analysis related to the topic of our work, and they were able to correlate signatures of criticality in the ongoing activity of barrel cortex of urethane anesthetized rats, depending on the cortical state, to an increase in the dynamic range. While in all the rats that we analyzed, we both found signatures of criticality (in terms of neuronal avalanches) and an increase in the power of the intrinsic IMF 2 rhythm, it would be an interesting future direction to correlate signatures of criticality to the relative amount of stimulus encoding, also depending on the level of anesthesia or on the cortical state.

Finally, the present work contributes to show that these ≈ 11 Hz spindle-like oscillations may be a relevant collective neural activity for the processing of whisker-related sensory input in the barrel cortex, and it is important to notice that their frequency is remarkably close to that of whisker twitching (7-12 Hz). The fact that our model admits a limit cycle in the barrel cortex has an important consequence in terms of oscillation theory. Indeed, in oscillation theory, the existence of a limit cycle or self-sustained oscillation is the basic ingredient for synchronization phenomena [63]. So, given the closeness of the frequency ranges of whisker (7-12 Hz) and barrel oscillations (≈ 11 Hz), we hypothesize that a synchronization mechanism could facilitate information transfer between these regions over a wide frequency range. Notably, we verify this hypothesis in a different work [82]. Our results thus support the hypothesis that spontaneous neural activity in the barrel cortex plays a significant physiological role, priming the barrel cortex for efficient processing of sensory input from whisker movements.

## 3 Methods

### Surgical preparation

All experimental procedures were approved by the University of Padova Animal Welfare Body (Organismo Preposto al Benessere Animale, OPBA) and by the Italian Ministry of Health, Directorate General for Animal Health and Veterinary Medicinal Products (authorization number 522/2018-PR). Wistar rats are kept in the animal research facility of the Department of Biomedical Sciences of the University of Padua under standard environmental conditions. Four young adult rats undergoing the experiments are in the range of P25-P35 (being P0 on the day of birth) with a body weight between 80 and 160 g and of both genders. Electrophysiological signals are acquired at 25 kHz from a 32 iridium-oxide electrodes array using an RHS stimulation/recording controller from Intan Technologies (Los Angeles, California, USA). Rats are anesthetized with an intraperitoneal induction dose of urethane (0.15/100 g of the body weight), followed after half an hour by an additional dose (0.015/100 g of the body weight) and after other 10 minutes by a sub-cutaneous dose of Carprofen painkiller (Rimadyl; 0.5 mg/100 g of the body weight). The animal is then positioned on a stereotaxic frame and the head is fixed by teeth- and ear-bars. Body temperature is maintained at 37°C by a heating pad and monitored by a rectal probe using a homeothermic monitoring system (World Precision Instruments ATC1000 DC temperature controller) throughout the procedure. An anterior-posterior opening in the skin is made in the center of the head and a dedicated window in the skull is drilled over the right somatosensory barrel cortex at stereotaxic coordinates from− 1 to −4 AP, from +4 to +8 ML referred to bregma [75, 81], and the probe is inserted orthogonally to the cortical surface using a micromanipulator (PatchStar; Scientifica). The depth is set at 0 when the electrode proximal to the chip tip touches the cortical surface. An Ag/AgCl electrode bathed in standard Krebs solution (in mM: NaCl 120, KCl 1.99, NaHCO 3 25.56, KH 2 PO 4 136.09, CaCl 2 2, MgSO 4 1.2, glucose 11) in proximity of the probe is used as a reference.

### Electrophysiological Recordings

Thalamocortical signals are recorded by inserting a 32-channels linear probe (E32+R-200-S1-L20 NT, Atlas Neuro, Leuven, Belgium, with a pitch between the electrodes of 200 *µm*) at coordinates − 2.9 AP, +6.4 ML. This position allows to acquire simultaneous signals from the barrel cortex and the ventral posteromedial nucleus (VPM) of the thalamus using 30 out of the 32 channels. Whiskers are cut to 10 mm and individually inserted up to 8 mm into a 25G hypodermic needle (BD Plastipak, Madrid, Spain), that is glued to a multilayer piezoelectric bender with integrated strain gauges and powered by a power amplifier (P-871.122 & E-650.00, Physik Instrumente, Karlsruhe, Germany). The mechanical stimulation is controlled by a waveform generator (Agilent 33250A 80 MHz, Agilent Technologies Inc., Colorado, USA), whose output is recorded synchronously with the neural signal. The most responsive whisker, i. e. the one providing the highest evoked LFP amplitude, is then selected for recording and stimulation. Data are collected in 5-minutes sets of 30 stimulation trials, while stimulations of the whisker are administered mechanically by a displacement of the cannula of 5 ms duration and 100 *µ*s rise/fall time every 10 seconds. This long time between stimuli ensures protection against any dynamic adaptation to stimuli [18]. In the thalamocortical recordings an artifact at 100 Hz is evident, and it is eliminated in the LFPs through a notch filter. To extract Local Field Potentials data from the raw recording, the signals are filtered straight and reverse to achieve linear phase with a second-order bandpass Butterworth IIR filter with a cutoff frequency of 1 Hz (lower) and 70 Hz (higher). To extract multiunits activity data (MUAs) from the raw recording, the signals are filtered straight and reverse to achieve linear phase with a second-order bandpass Butterworth IIR filter with a cutoff frequency of 300 Hz (lower) and 3 kHz (higher). The spatial average of the signal recorded by the 32 channels is removed from each channel to reduce the correlated noise [70].

### Histological verification of probe insertion

The position of the recording probe within S1 cortex and thalamus estimated by LFP profiles [55, 83] was verified histologically in sample experiments. To this purpose, at the end of the recording session the rat was sacrificed and, after careful withdrawal of the probe, the brain explanted, formalin fixed and paraffin-embedded. Five micrometers thick coronal slices were stained by Nissl cresyl violet (see Supplementary Materials)

### Data analysis

#### Firing Rate

MUAs are recognized by detecting negative peaks below a threshold of 4/0.675 times the median of the absolute value of the signal [70]. The firing rate is estimated by the sum in a moving window of 60 ms of the spikes detected in each channel.

#### Empirical Mode Decomposition (EMD) and intrinsic mode functions (IMF)

We use empirical mode decomposition (EMD) [39] through the EMD package [69] in order to study the frequency content of non-stationary and non-linear physiological signals. The EMD isolates a small number of temporally adaptive basis functions (intrinsic mode functions, IMF) and derives the dynamics in frequency and amplitude directly from them. The IMFs are obtained through an iterative process called sifting [39], that ensures that the EMDs are symmetric with respect to the local zero mean, and have the same numbers of zero crossings and extrema. This ensures a well-defined instantaneous frequency. Indeed, after performing the Hilbert transform [84] on each IMF component *Y*_*j*_, and after computing the instantaneous frequency as 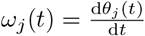 (where 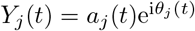), we can express the data in the following form

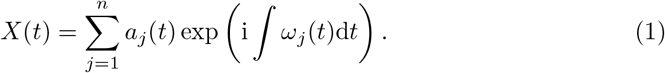

where the index *j* indicates the j-th IMF and *ω*_*j*_(*t*) and *a*_*j*_(*t*) are respectively the instantaneous frequency and amplitude for IMF j at time t. Thus, we have broken through the restriction of the constant amplitude and fixed-frequency Fourier expansion, and arrived at a variable amplitude and frequency representation.

Equation 1 also enables us to represent the amplitude and the instantaneous frequency as functions of time in a three-dimensional plot, in which the amplitude can be contoured on the frequency-time plane. This frequency-time distribution of the amplitude is designated as the Hilbert amplitude spectrum, *H*(*ω, t*), or simply Hilbert spectrum. In particular, in this work, we have considered the marginal Hilbert Spectrum,

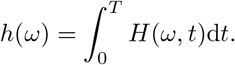

The marginal spectrum offers a measure of total amplitude (or energy) contribution from each frequency value. The frequency in either *H*(*ω*; *t*) or *h*(*ω*) has a totally different meaning from the Fourier spectral analysis. In the Fourier representation, the existence of energy at a frequency, *ω*, means a component of a sine or a cosine wave persisted. through the time span of the data. Here, the existence of energy at the frequency, *ω*, means only that, in the whole time span of the data, there is a higher likelihood for such a wave to have appeared locally. In fact, the Hilbert spectrum is a weighted non-normalized joint amplitude-frequency-time distribution.

To a correct application of the EMD algorithm, common practice states that the IMFs should be well separated - the components should have low correlations/orthogonality scores - and the instantaneous frequency content of the IMFs should not strongly overlap (no mode mixing). This can be achieved by using a pseudo-mode splitting index (PMSI) [23]. In particular, we applied to the local field potential recordings a mask-sift algorithm, that overcomes some limitations of the standard sift algorithm with intermittent and noisy signals [16], by masking some chosen frequencies in the signals so to avoid mode mixing. Indeed, choosing a frequency mask effectively puts a lower bound on the frequency content that can enter a particular IMF. The frequency of the first mask was set to 30 Hz, and the mask step factor was set to 2 (default value) [69] (by simulations, it was shown that a mask at a given frequency *f* suppresses frequencies below 0.7 × *f* [29]) (see Supplementary Materials for other details of the EMD analysis).

#### Phase-amplitude coupling, phase-avalanche coupling

Phase amplitude coupling is computed by using the EMD package [69] between the second IMF amplitude and the fourth IMF phase. Cycles are detected in the fourth IMF, and the corresponding phase is binned. Then, the amplitude of IMF 2 is associated with its respective phase bin, and the amplitude values are averaged across all the cycles that are found. The same procedure is applied to study the coupling between the phase of IMF 4 and the fraction of avalanches. The only difference is that the binned phases of IMF 4 are associated with the corresponding fraction-of-avalanches value.

#### Avalanches’ statistics and avalanches’ density

Avalanches are computed using standardized pipelines [4, 53]. MUAs are recognized by detecting negative peaks below a threshold of 3/0.675 times the median of the absolute value of the signal. Then, the average inter-event interval is computed [4] to bin the data. Indeed, the distributions of inter-event intervals that we found, both in the barrel cortex and in the thalamus, were not heavy-tailed, nor bimodal, hence we employed the standard practice to take the mean of the distribution as the reference time bin for the avalanches. This resulted in the following ISI for avalanches analysis in the ongoing activity of the barrel cortex: 5.37 ± 0.04 ms, 3.57 ± 0.03 ms, 4.04 ± 0.03 ms, 4.12 ± 0.03 ms; and the following ones for the thalamus: 2.74 ± 0.01 ms, 2.60 ± 0.01 ms, 1.97 ± 0.01 ms, 1.66 ± 0.004 ms. Avalanches are then detected as sequences of time bins with activity, separated by empty bins. The number of events in an avalanche is the size of an avalanche, while the number of bins is the duration of an avalanche. They are fitted with power-laws using a corrected maximum-likelihood method [53]. Indeed, the avalanches sizes and durations distributions are fitted using the maximum likelihood method. The fitting function for both avalanche sizes and duration is a discrete power-law:

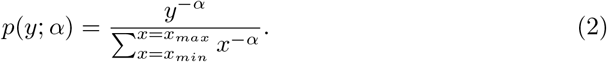

The parameter *x*_*max*_ is set to the maximum observed size or duration. Then the tails of the distributions are fitted by selecting as parameter *x*_*min*_ the one that minimizes the Kolmogorov-Smirnov distance (KS), following the method proposed by Clauset et al. [12]:

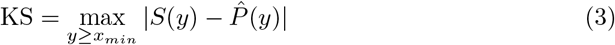

where *S*(*y*) is the cumulative distribution function (CDF) of the data and 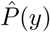 is the CDF of the theoretical distribution fitted with the parameter that best fits the data for *y* ≥ *x*_*min*_.

After finding the best-fit power law, to assess goodness-of-fit we compared the experimental data against 1000 surrogate datasets drawn from the best-fit power law distribution with the same number of samples as the experimental dataset. The deviation between the surrogate datasets and a perfect power law was quantified with the KS statistic. The p-value of the power-law fit was defined as the fraction of these surrogate KS statistics which were greater than the KS statistic for the experimental data. Note that the data were considered power law distributed if the null hypothesis could not be rejected, namely if the the p-value turned out to be greater than the significance level, which was set to a conservative value of 0.1.

Then, the avalanche density as a function of time is calculated, as it was proposed in [72], i.e., it is computed as the fraction of time occupied by avalanches in a time *T*_0_, with a sliding step of 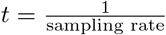. *T*_0_ was set to 5 × average inter event interval.

## Reduced model of the barrel cortex

Brain activity in the barrel cortex was modeled using a set of equations similar to the well-known Wilson-Cowan system [91]. This model has been previously used to describe the barrel cortex of rats, providing a good qualitative and quantitative agreement with anatomical and physiological properties of the barrel cortex as well as its response to whisker stimulation [64]. This model is the result of a reduction of a biologically detailed spike-model of a single barrel previously developed by Kyriazi and Simons [46] (with 70 excitatory and 30 inhibitory neurons, modeled as leaky linear integrators). The resulting model consists of one excitatory and one inhibitory neuronal population simulating the barrel cortex, connected with each other and receiving thalamic input. The thalamic input is introduced in the form of time histograms (TH). The corresponding equations describe the dynamics of the average synaptic drive of both excitatory and inhibitory populations,

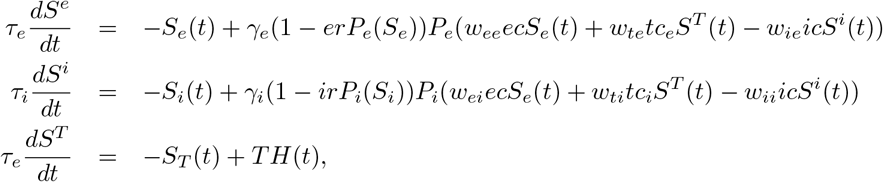

The reduced equations succinctly define the relationship between three distinct measures of neuronal activity (see equation 4, in which for simplicity we don’t consider the refractory effect), as mentioned in the main text: voltage, firing rate, and a measure which we describe as synaptic drive (*S*_*e*_ and *S*_*i*_ are the synaptic drives and are linked to the firing rate through the conversion factors *γ*_*e*_, *γ*_*i*_ of the excitatory and inhibitory populations, respectively).

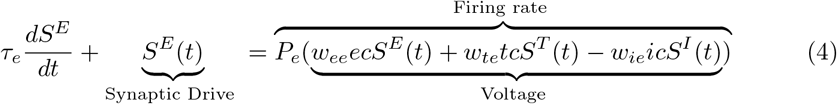

*P*_*e*_ and *P*_*i*_ are activation functions (excitatory and inhibitory), given by,

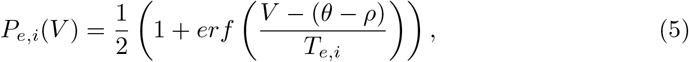

where *V* is the membrane potential and *T*_*e,i*_ is the temperature for activation function of the excitatory and inhibitory populations. The parameters *θ* and *ρ* are the firing threshold and the resting membrane potential, respectively. Here *erf* (*x*) is the Gauss error function, defined in the usual way,

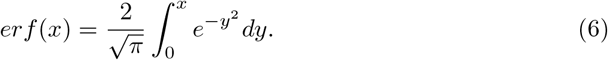

The parameters of the model used in this work are summarized in table 2 and were calibrated experimentally [46].

In our case, we give as input to the model the time histograms of thalamus activity either after stimulation of the rat whisker or during spontaneous activity. Note that the balance of feedforward (thalamic) inhibition over excitation 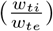 and the refractory parameter *er* are varied (see Table 2, but within the standard deviations provided by [46]) so that the model can admit a limit cycle and behaves as an amplifier, as predicted in [65] (while it behaves as a damper with higher values of *w*_*ti*_).

### Avalanches analysis in the model

Avalanches and the coupling of the fraction of avalanches with the slow oscillation are measured in the model in the same way as they are analyzed in the data. For this purpose, spiking events from a number of artificial neurons of the same order of magnitude as the number of channels that we have, is generated. To generate the spiking data, each neuron is modeled as an inhomogeneous Poisson process with a time-varying rate equal to the firing rate of the cortical population [48]. Note that with this choice, the population firing rate equals the firing rate generated by the model. Then, the average inter-event interval is computed and avalanches are studied in the same way as in the data.

## 4 Author contributions

S.S. and S.V. conceived the study and were in charge of overall direction and planning. M.M., and A.L. carried out the experiments. R.G., B.M., and S.S. designed the theoretical framework. B. M. analyzed the data and M. T. provided guidance in the data analysis. B.M. and R.G. planned the simulations and B. M. carried out and analyzed the simulations. All authors contributed to the interpretation of the results, provided critical feedback, and helped shape the research, analysis, and manuscript.

## 5 Acknowledgments

S.S. acknowledges NEXTGENERATIONEU (NGEU) and is funded by the Ministry of University and Research (MUR), National Recovery and Resilience Plan (NRRP), project MNESYS (PE0000006) – A Multiscale integrated approach to the study of the nervous system in health and disease (DN. 1553 11.10.2022). B.M. acknowledges the DFA UNIPD for funding through the PARD 2023 project “Response theory for brain network discovery and control”. The funders had no role in study design, data collection and analysis, decision to publish, or preparation of the manuscript. The authors are grateful to Dr. Aron Emmi for help with histology.

## Supporting Information

**A Sec**. IMFs selection

**S1 Fig**. Phase diagram of the model

**S2 Fig**. Intrinsic Mode functions

**S3 Fig**. Relationship between the fast oscillation and avalanches.

**S4 Fig**. Marginal Hilbert spectrum of intrinsic Mode Functions 2 and 4 in each of the four rats analyzed (1, 2, 3, 4). (1) (a) Marginal Hilbert spectrum after the whisker stimulation in the rat barrel cortex. (b) Marginal Hilbert spectrum during spontaneous activity in the rat barrel cortex. (c) Marginal Hilbert spectrum after the whisker stimulation in the rat thalamus. (d) Marginal Hilbert spectrum during spontaneous activity in the rat thalamus. Same in 2, 3, 4, for three additional rats.

**S5 Fig**. Avalanches distribution in the barrel cortex during spontaneous activity (1) and after stimulation of the whisker (2) Avalanches distribution in the thalamus during spontaneous activity (3) and after stimulation of the whisker (4) (Rat 1)

**S6 Fig**. Avalanche distributions for an additional rat (rat 2), same analysis as in Figure S5

**S7 Fig**. Avalanche distributions for an additional rat (rat 3), same analysis as in Figure S5

**S8 Fig**. Avalanche distributions for an additional rat (rat 4), same analysis as in Figure S5

**S9 Fig**. Phase amplitude coupling (PAC) between the phase of Intrinsic mode function 4 and the amplitude of intrinsic mode function 2

**S10 Fig**. Coupling between the phase of Intrinsic mode function 4 and the fraction of avalanches

**S11 Fig**. Histological verification of probe insertion and LFPs profile

**S12 Fig**. Results for the model (with Rat 1 firing rate as input)

**S13 Fig**. Results for the model (with Rat 2 firing rate as input). Same as in S12, for rat 2.

**S14 Fig**. Results for the model (with Rat 3 firing rate as input). Same as in S12, for rat 3.

**S15 Fig**. Results for the model (with Rat 4 firing rate as input). Same as in S12, for rat 4.

## References

1. Y. Ahmadian and K. D. Miller. What is the dynamical regime of cerebral cortex? Neuron, 109(21):3373–3391, 2021.

2. G. Barzon, G. Nicoletti, B. Mariani, M. Formentin, and S. Suweis. Criticality and network structure drive emergent oscillations in a stochastic whole-brain model. Journal of Physics: Complexity, 3(2):025010, 2022.

3. D. Battaglia and D. Hansel. Synchronous chaos and broad band gamma rhythm in a minimal multi-layer model of primary visual cortex. PLoS computational biology, 7(10):e1002176, 2011.

4. J. M. Beggs and D. Plenz. Neuronal avalanches in neocortical circuits. Journal of Neuroscience, 12:23(35):11167–11177, 2003.

5. M. Bonnefond, S. Kastner, and O. Jensen. Communication between brain areas based on nested oscillations. eneuro, 4(2), 2017.

6. V. Buendía, P. Villegas, R. Burioni, and M. A. Muñoz. Hybrid-type synchronization transitions: Where incipient oscillations, scale-free avalanches, and bistability live together. Phys. Rev. Res., 3:023224, Jun 2021.

7. G. Buzsaki. Rhythms of the Brain. Oxford University Press, 2006.

8. G. Buzsáki, N. Logothetis, and W. Singer. Scaling brain size, keeping timing: evolutionary preservation of brain rhythms. Neuron, 80(3):751–764, 2013.

9. E. Capek, T. L. Ribeiro, P. Kells, K. Srinivasan, S. R. Miller, E. Geist, M. Victor, A. Vakili, S. Pajevic, D. R. Chialvo, et al. Parabolic avalanche scaling in the synchronization of cortical cell assemblies. Nature communications, 14(1):2555, 2023.

10. R. L. Carhart-Harris, R. Leech, P. J. Hellyer, M. Shanahan, A. Feilding, E. Tagliazucchi, D. R. Chialvo, and D. Nutt. The entropic brain: a theory of conscious states informed by neuroimaging research with psychedelic drugs. Frontiers in human neuroscience, 8:55875, 2014.

11. D. R. Chialvo. Emergent complex neural dynamics. Nature physics, 6(10):744–750, 2010.

12. A. Clauset, C. R. Shalizi, and M. E. Newman. Power-law distributions in empirical data. SIAM review, 51(4):661–703, 2009.

13. M. X. Cohen. Analyzing neural time series data: theory and practice. MIT press, 2014.

14. C. Curto, S. Sakata, S. Marguet, V. Itskov, and K. D. Harris. A simple model of cortical dynamics explains variability and state dependence of sensory responses in urethane-anesthetized auditory cortex. Journal of Neuroscience, 29(34):10600–10612, 2009.

15. G. Deco, V. K. Jirsa, and A. R. McIntosh. Emerging concepts for the dynamical organization of resting-state activity in the brain. Nature reviews neuroscience, 12(1):43–56, 2011.

16. R. Deering and J. F. Kaiser. The use of a masking signal to improve empirical mode decomposition. In Proceedings.(ICASSP’05). IEEE International Conference on Acoustics, Speech, and Signal Processing, 2005., volume 4, pages iv–485. IEEE, 2005.

17. D. Derdikman, R. Hildesheim, E. Ahissar, A. Arieli, and A. Grinvald. Imaging spatiotemporal dynamics of surround inhibition in the barrels somatosensory cortex. Journal of Neuroscience, 23(8):3100–3105, 2003.

18. D. Derdikman, C. Yu, S. Haidarliu, K. Bagdasarian, A. Arieli, and E. Ahissar. Layer-specific touch-dependent facilitation and depression in the somatosensory cortex during active whisking. Journal of Neuroscience, 26(37):9538–9547, 2006.

19. S. Di Santo, P. Villegas, R. Burioni, and M. A. Muñoz. Landau–ginzburg theory of cortex dynamics: Scale-free avalanches emerge at the edge of synchronization. Proceedings of the National Academy of Sciences, 115(7):E1356–E1365, 2018.

20. M. E. Diamond, M. Von Heimendahl, P. M. Knutsen, D. Kleinfeld, and E. Ahissar. ‘where’and’what’in the whisker sensorimotor system. Nature Reviews Neuroscience, 9(8):601–612, 2008.

21. A. Dimakou, G. Pezzulo, A. Zangrossi, and M. Corbetta. The predictive nature of spontaneous brain activity across scales and species. Neuron, 113(9):1310–1332, 2025.

22. V. Douchamps, M. Di Volo, A. Torcini, D. Battaglia, and R. Goutagny. Gamma oscillatory complexity conveys behavioral information in hippocampal networks. Nature Communications, 15(1):1849, 2024.

23. M. S. Fabus, A. J. Quinn, C. E. Warnaby, and M. W. Woolrich. Automatic decomposition of electrophysiological data into distinct nonsinusoidal oscillatory modes. Journal of Neurophysiology, 126(5):1670–1684, 2021. PMID: 34614377.

24. E. E. Fanselow and M. A. Nicolelis. Behavioral modulation of tactile responses in the rat somatosensory system. Journal of neuroscience, 19(17):7603–7616, 1999.

25. E. E. Fanselow, K. Sameshima, L. A. Baccala, and M. A. Nicolelis. Thalamic bursting in rats during different awake behavioral states. Proceedings of the National Academy of Sciences, 98(26):15330–15335, 2001.

26. A. J. Fontenele, N. A. P. de Vasconcelos, T. Feliciano, L. A. A. Aguiar, C. Soares-Cunha, B. Coimbra, L. Dalla Porta, S. Ribeiro, A. J. a. Rodrigues, N. Sousa, P. V. Carelli, and M. Copelli. Criticality between cortical states. Phys. Rev. Lett., 122:208101, 2019.

27. A. J. Fontenele, J. S. Sooter, V. K. Norman, S. H. Gautam, and W. L. Shew. Low-dimensional criticality embedded in high-dimensional awake brain dynamics. Science Advances, 10(17):eadj9303, 2024.

28. L. Fontolan, B. Morillon, C. Liegeois-Chauvel, and A.-L. Giraud. The contribution of frequency-specific activity to hierarchical information processing in the human auditory cortex. Nature communications, 5(1):4694, 2014.

29. O. B. Fosso and M. Molinas. Method for mode mixing separation in empirical mode decomposition. arXiv preprint arXiv:1709.05547, 2017.

30. D. Fraiman and D. R. Chialvo. What kind of noise is brain noise: anomalous scaling behavior of the resting brain activity fluctuations. Frontiers in physiology, 3:307, 2012.

31. N. Friedman, S. Ito, B. A. Brinkman, M. Shimono, R. L. DeVille, K. A. Dahmen, J. M. Beggs, and T. C. Butler. Universal critical dynamics in high resolution neuronal avalanche data. Physical review letters, 108(20):208102, 2012.

32. P. Fries. A mechanism for cognitive dynamics: neuronal communication through neuronal coherence. Trends in cognitive sciences, 9(10):474–480, 2005.

33. S. H. Gautam, T. T. Hoang, K. McClanahan, S. K. Grady, and W. L. Shew. Maximizing sensory dynamic range by tuning the cortical state to criticality. PLOS Computational Biology, 11(12):1–15, 12 2015.

34. E. D. Gireesh and D. Plenz. Neuronal avalanches organize as nested theta-and beta/gamma-oscillations during development of cortical layer 2/3. Proceedings of the National Academy of Sciences, 105(21):7576–7581, 2008.

35. J. J. Harris, R. Jolivet, and D. Attwell. Synaptic energy use and supply. Neuron, 75(5):762–777, 2012.

36. R. F. Helfrich, B. A. Mander, W. J. Jagust, R. T. Knight, and M. P. Walker. Old brains come uncoupled in sleep: Slow wave-spindle synchrony, brain atrophy, and forgetting. Neuron, 97, 2018.

37. K. B. Hengen and W. L. Shew. Is criticality a unified setpoint of brain function? Neuron, 113(16):2582–2598.e2, 2025.

38. J. Hidalgo, J. Grilli, S. Suweis, M. A. Munoz, J. R. Banavar, and A. Maritan. Information-based fitness and the emergence of criticality in living systems. Proceedings of the National Academy of Sciences, 111(28):10095–10100, 2014.

39. N. E. Huang, Z. Shen, S. R. Long, M. C. Wu, H. H. Shih, Q. Zheng, N.-C. Yen, C. C. Tung, and H. H. Liu. The empirical mode decomposition and the hilbert spectrum for nonlinear and non-stationary time series analysis. Proceedings of the Royal Society of London. Series A: mathematical, physical and engineering sciences, 454(1971):903–995, 1998.

40. A. Hyafil, A.-L. Giraud, L. Fontolan, and B. Gutkin. Neural cross-frequency coupling: connecting architectures, mechanisms, and functions. Trends in neurosciences, 38(11):725–740, 2015.

41. J. Ito, S. Roy, Y. Liu, Y. Cao, M. Fletcher, L. Lu, J. Boughter, S. Grün, and D. Heck. Whisker barrel cortex delta oscillations and gamma power in the awake mouse are linked to respiration. Nature communications, 5(1):3572, 2014.

42. S. A. Jones, J. H. Barfield, V. K. Norman, and W. L. Shew. Scale-free behavioral dynamics directly linked with scale-free cortical dynamics. eLife, 12:e79950, jan 2023.

43. R. Khazipov, A. Sirota, X. Leinekugel, G. L. Holmes, Y. Ben-Ari, and G. Buzsáki. Early motor activity drives spindle bursts in the developing somatosensory cortex. Nature, 432(7018):758–761, 2004.

44. D. Kleinfeld, E. Ahissar, and M. E. Diamond. Active sensation: insights from the rodent vibrissa sensorimotor system. Current opinion in neurobiology, 16(4):435–444, 2006.

45. N. Kopell. We got rhythm: Dynamical systems of the nervous system. Notices of the AMS, 47(1):6–16, 2000.

46. H. T. Kyriazi and D. J. Simons. Thalamocortical response transformations in simulated whisker barrels. Journal of Neuroscience, 13(4):1601–1615, 1993.

47. P. Lakatos, A. S. Shah, K. H. Knuth, I. Ulbert, G. Karmos, and C. E. Schroeder. An oscillatory hierarchy controlling neuronal excitability and stimulus processing in the auditory cortex. Journal of neurophysiology, 94(3):1904–1911, 2005.

48. P. W. Lewis and G. S. Shedler. Simulation of nonhomogeneous poisson processes by thinning. Naval research logistics quarterly, 26(3):403–413, 1979.

49. F. Lombardi, H. J. Herrmann, D. Plenz, and L. De Arcangelis. On the temporal organization of neuronal avalanches. Frontiers in systems neuroscience, 8:204, 2014.

50. F. Lombardi, S. Pepić, O. Shriki, G. Tkačik, and D. De Martino. Statistical modeling of adaptive neural networks explains co-existence of avalanches and oscillations in resting human brain. Nature Computational Science, 3, 2023.

51. M. Malekmohammadi, W. J. Elias, and N. Pouratian. Human Thalamus Regulates Cortical Activity via Spatially Specific and Structurally Constrained Phase-Amplitude Coupling. Cerebral Cortex, 25(6):1618–1628, 01 2014.

52. D. Mantini, M. G. Perrucci, C. Del Gratta, G. L. Romani, and M. Corbetta. Electrophysiological signatures of resting state networks in the human brain. Proceedings of the National Academy of Sciences, 104(32):13170–13175, 2007.

53. B. Mariani, G. Nicoletti, M. Bisio, M. Maschietto, R. Oboe, A. Leparulo, S. Suweis, and S. Vassanelli. Neuronal avalanches across the rat somatosensory barrel cortex and the effect of single whisker stimulation. Frontiers in Systems Neuroscience, 15:89, 2021.

54. B. Mariani, G. Nicoletti, M. Bisio, M. Maschietto, S. Vassanelli, and S. Suweis. Disentangling the critical signatures of neural activity. Scientific reports, 12(1):10770, 2022.

55. H. S. Meyer, R. Egger, J. M. Guest, R. Foerster, S. Reissl, and M. Oberlaender. Cellular organization of cortical barrel columns is whisker-specific. Proceedings of the national academy of sciences, 110(47):19113–19118, 2013.

56. S. R. Miller, S. Yu, and D. Plenz. The scale-invariant, temporal profile of neuronal avalanches in relation to cortical *γ*–oscillations. Scientific reports, 9(1):16403, 2019.

57. M. A. Munoz. Colloquium: Criticality and dynamical scaling in living systems. Reviews of Modern Physics, 90(3):031001, 2018.

58. G. T. Neske. The slow oscillation in cortical and thalamic networks: mechanisms and functions. Frontiers in neural circuits, 9:88, 2016.

59. J. O’Byrne and K. Jerbi. How critical is brain criticality? Trends in neurosciences, 45(11):820–837, 2022.

60. T. Petermann, T. C. Thiagarajan, M. A. Lebedev, M. A. Nicolelis, D. R. Chialvo, and D. Plenz. Spontaneous cortical activity in awake monkeys composed of neuronal avalanches. Proceedings of the National Academy of Sciences, 106(37):15921–15926, 2009.

61. C. C. Petersen. The functional organization of the barrel cortex. Neuron, 56(2):339–355, 2007.

62. G. Pezzulo, M. Zorzi, and M. Corbetta. The secret life of predictive brains: what’s spontaneous activity for? Trends in cognitive sciences, 25(9):730–743, 2021.

63. A. Pikovsky, M. Rosenblum, and J. Kurths. Synchronization: a universal concept in nonlinear science, 2002.

64. D. J. Pinto, J. C. Brumberg, D. J. Simons, G. B. Ermentrout, and R. Traub. A quantitative population model of whisker barrels: re-examining the wilson-cowan equations. Journal of computational neuroscience, 3:247–264, 1996.

65. D. J. Pinto, J. A. Hartings, J. C. Brumberg, and D. J. Simons. Cortical damping: analysis of thalamocortical response transformations in rodent barrel cortex. Cerebral cortex, 13(1):33–44, 2003.

66. S.-S. Poil, R. Hardstone, H. D. Mansvelder, and K. Linkenkaer-Hansen. Critical-state dynamics of avalanches and oscillations jointly emerge from balanced excitation/inhibition in neuronal networks. Journal of Neuroscience, 32(29):9817–9823, 2012.

67. A. Ponce-Alvarez, G. Deco, P. Hagmann, G. L. Romani, D. Mantini, and M. Corbetta. Resting-state temporal synchronization networks emerge from connectivity topology and heterogeneity. PLoS computational biology, 11(2):e1004100, 2015.

68. A. Ponce-Alvarez, A. Jouary, M. Privat, G. Deco, and G. Sumbre. Whole-brain neuronal activity displays crackling noise dynamics. Neuron, 100(6):1446–1459.e6, 2018.

69. A. J. Quinn, V. Lopes-dos Santos, D. Dupret, A. C. Nobre, and M. W. Woolrich. Emd: Empirical mode decomposition and hilbert-huang spectral analyses in python. Journal of Open Source Software, 6(59):2977, 2021.

70. R. Q. Quiroga, Z. Nadasdy, and Y. Ben-Shaul. Unsupervised Spike Detection and Sorting with Wavelets and Superparamagnetic Clustering. Neural Computation, 16(8):1661–1687, 08 2004.

71. M. E. Raichle. The restless brain. Brain connectivity, 1(1):3–12, 2011.

72. S. Scarpetta, N. Morisi, C. Mutti, N. Azzi, I. Trippi, R. Ciliento, I. Apicella, G. Messuti, M. Angiolelli, F. Lombardi, et al. Criticality of neuronal avalanches in human sleep and their relationship with sleep macro-and micro-architecture. Iscience, 26(10), 2023.

73. K. Semba, H. Szechtman, and B. R. Komisaruk. Synchrony among rhythmical facial tremor, neocortical ‘alpha’waves, and thalamic non-sensory neuronal bursts in intact awake rats. Brain research, 195(2):281–298, 1980.

74. F.-Z. Shaw. Is spontaneous high-voltage rhythmic spike discharge in long evans rats an absence-like seizure activity? Journal of neurophysiology, 91(1):63–77, 2004.

75. S. Shimegi, T. Akasaki, T. Ichikawa, and H. Sato. Physiological and anatomical organization of multiwhisker response interactions in the barrel cortex of rats. J. Neurosci., 20(16):6241–6248, Aug. 2000.

76. W. Singer. Neuronal oscillations: unavoidable and useful? European Journal of Neuroscience, 48(7):2389–2398, 2018.

77. S. M. Smith, P. T. Fox, K. L. Miller, D. C. Glahn, P. M. Fox, C. E. Mackay, N. Filippini, K. E. Watkins, R. Toro, A. R. Laird, et al. Correspondence of the brain’s functional architecture during activation and rest. Proceedings of the national academy of sciences, 106(31):13040–13045, 2009.

78. A. Sobolewski, D. A. Swiejkowski, A. Wróbel, and E. Kublik. The 5–12 hz oscillations in the barrel cortex of awake rats–sustained attention during behavioral idling? Clinical Neurophysiology, 122(3):483–489, 2011.

79. V. Sorrenti, C. Cecchetto, M. Maschietto, S. Fortinguerra, A. Buriani, and S. Vassanelli. Understanding the effects of anesthesia on cortical electrophysiological recordings: A scoping review. International Journal of Molecular Sciences, 22(3), 2021.

80. C. Stringer, M. Pachitariu, N. Steinmetz, C. B. Reddy, M. Carandini, and K. D. Harris. Spontaneous behaviors drive multidimensional, brainwide activity. Science, 364(6437):eaav7893, 2019.

81. L. W. Swanson. Brain maps 4.0-structure of the rat brain: An open access atlas with global nervous system nomenclature ontology and flatmaps. J. Comp. Neurol., 526(6):935–943, Apr. 2018.

82. M. Tambaro, R. Guevara, M. Maschietto, A. L. Leparulo, C. Cecchetto, G. Nicoletti, B. Mariani, S. Suweis, and V. Stefano. The somatosensory barrel cortex controls the spindle thalamocortical oscillation by frequency locking. biorXiv, 2025.

83. S. Temereanca and D. J. Simons. Local field potentials and the encoding of whisker deflections by population firing synchrony in thalamic barreloids. Journal of neurophysiology, 89(4):2137–2145, 2003.

84. E. C. Titchmarsh. Introduction to the theory of Fourier integral. The Clarendon Press, 1937.

85. J. Touboul and A. Destexhe. Can power-law scaling and neuronal avalanches arise from stochastic dynamics? PLOS ONE, 5(2):1–14, 02 2010.

86. S. Venkatraman and J. M. Carmena. Behavioral modulation of stimulus-evoked oscillations in barrel cortex of alert rats. Frontiers in integrative neuroscience, page 10, 2009.

87. P. Villegas, S. di Santo, R. Burioni, and M. A. Muñoz. Time-series thresholding and the definition of avalanche size. Phys. Rev. E, 100:012133, Jul 2019.

88. J. L. Vincent, G. H. Patel, M. D. Fox, A. Z. Snyder, J. T. Baker, D. C. Van Essen, J. M. Zempel, L. H. Snyder, M. Corbetta, and M. E. Raichle. Intrinsic functional architecture in the anaesthetized monkey brain. Nature, 447(7140):83–86, 2007.

89. S. B. Vincent. The Functions of the Vibrissae in the Behavior of the White Rat…, volume 1. University of Chicago, 1912.

90. T. Volpi, E. Silvestri, M. Aiello, J. J. Lee, A. G. Vlassenko, M. S. Goyal, M. Corbetta, and A. Bertoldo. The brain’s “dark energy” puzzle: How strongly is glucose metabolism linked to resting-state brain activity? Journal of Cerebral Blood Flow & Metabolism, page 0271678X241237974, 2024.

91. H. R. Wilson and J. D. Cowan. Excitatory and inhibitory interactions in localized populations of model neurons. Biophysical journal, 12(1):1–24, 1972.

92. T. A. Woolsey and H. Van der Loos. The structural organization of layer iv in the somatosensory region (si) of mouse cerebral cortex: the description of a cortical field composed of discrete cytoarchitectonic units. Brain research, 17(2):205–242, 1970.

