## Supplemental Figures for "Whisker stimulation reinforces a resting-state network in the barrel cortex: nested oscillations and avalanches"

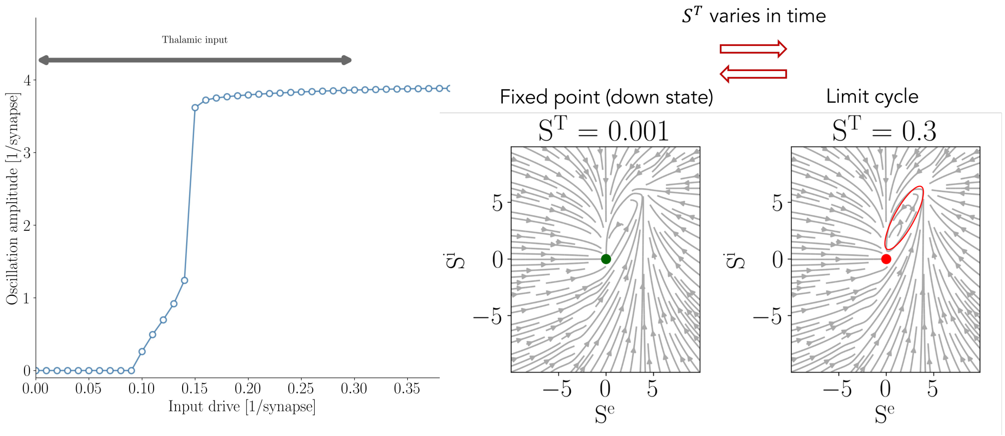


(1) (2)

FIG. S1: **Phase diagram of the model**. Panel (1): phase diagram of the cortical dynamics, indicating the amplitude of the oscillation of the (excitatory) cortical population, as a function of the thalamic input drive. It is possible to see that the oscillation arises above a threshold of $\approx 0.15$ [1/synapse]. Importantly, the input drive is not constant in our model, yet it is described by the time-varying experimental thalamic dynamics. Panel (2): the same features can be observed in the phase space, where above the previously mentioned threshold, a limit cycle can be appreciated in the streamlines of the dynamics.

### A. IMFs selection

The selection of IMFs was guided by their spectral features and temporal persistence, rather than by a posteriori selection. The first IMF is predominantly characterized by high-frequency noise, consistent with the well-established practice in EMD analysis of low pass filtering the data prior to analysis [1, 2]; indeed, the first mode captures residual high-frequency components, with a broad frequency content and no clear peak (as can be seen in the Hilbert spectrum in Figure S2 (a) and (b), where IMF 1 is colored in dark green), hence we did not consider it. The third IMF was not systematically analyzed because it does not exhibit a clear spectral peak in the power spectrum, and is temporally particularly confined. In figure (c) it is possible to observe the time evolution of the different IMFs, and in figure (d) the corresponding time-frequency analysis (based on Morlet wavelet), as a comparison. Indeed, IMF 3 brief bursts (highlighted by a shaded pink area in Figure S2 (c)) often precedes stimulus onset, suggesting that it reflects transient components (likely event-related components) rather than sustained oscillatory activity. Future analyses may investigate the interplay between IMF 3 and IMF 2, and whether IMF 3 contains relevant physiological frequencies, or just event-related components. The fourth IMF was selected based on its clear correspondence with the slow oscillation peak ($< 4$ Hz) (see Figure S2 (a) and Fig. (b)) observed in the power spectrum, representing a stable low-frequency component that persists throughout the recording. The later IMFs are characterized by progressively slower oscillations persisting throughout the entire recording, while the final IMF represents the residual of the decomposition (IMF 6) — that is, what remains once no further oscillatory modes can be extracted by the sifting process, conceptually analogous to the slow trend or baseline of the signal. The original signal can indeed be reconstructed by summing all extracted IMFs and the residual.

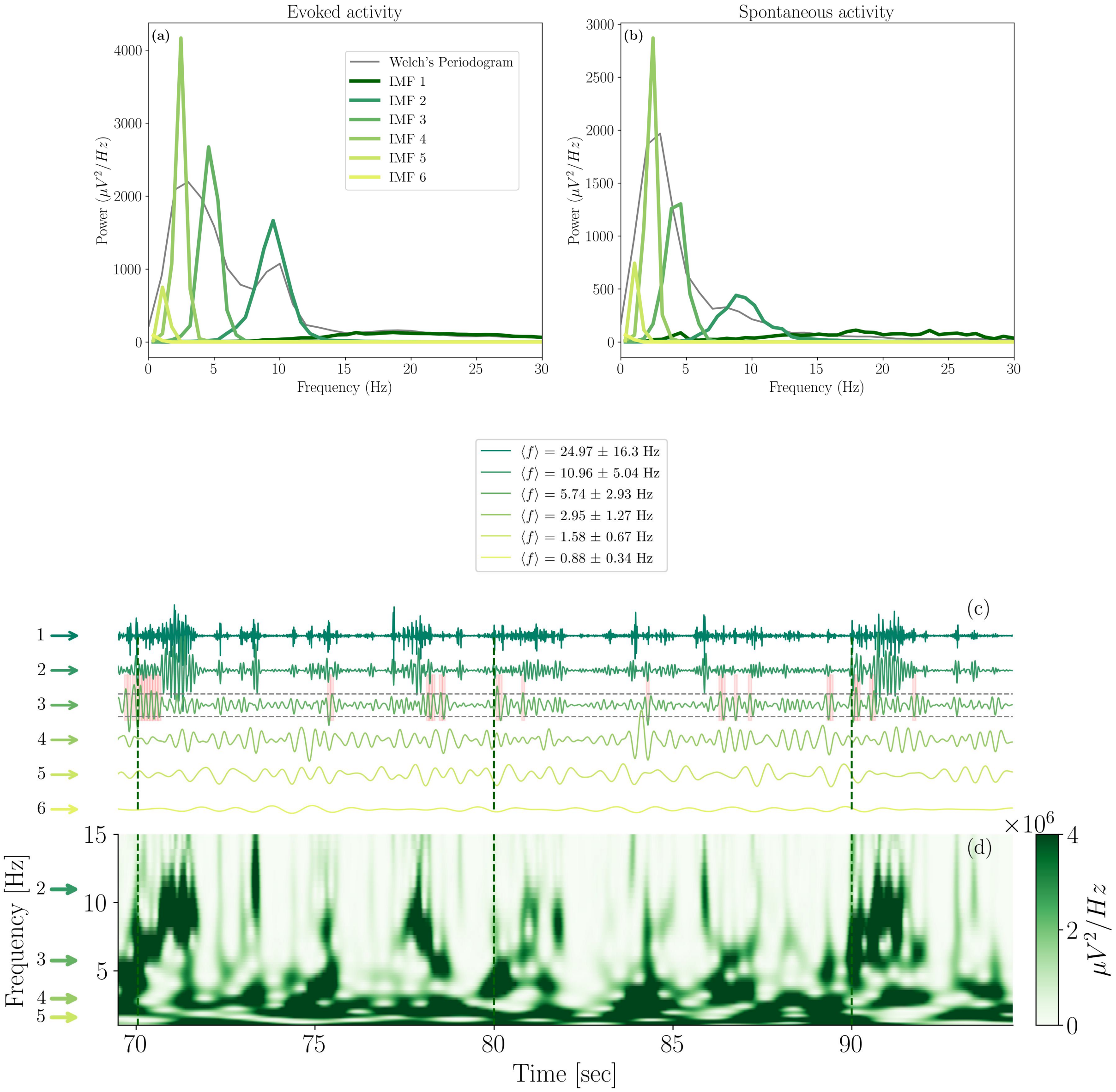


FIG. S2: **Intrinsic Mode functions.** Panels (a) and (b). The welch periodogram (standard power-spectrum) and the Hilbert spectrum of the 6 IMFs are reported in the periods of evoked activity (a) and spontaneous activity (b). These plots refer to all the trials of a rat, concatenated together. (c) Time evolution of the 6 intrinsic mode functions. Three stimulations of the whisker are highlighted by the vertical dashed line. (d) Time-frequency analysis based on Morlet wavelets to show the correspondence between a more standard Fourier method and the IMFs' dynamics. IMF number 1 is high-frequency noise and is not indicated by an arrow, as well as IMF number 6, which is the final residual.

Importantly, combining IMF 4 and IMF 5 into a single component yields results fully consistent with those reported in the manuscript, demonstrating that our findings are robust with respect to the precise separation of slow modes.

The IMF are calculated separately during evoked activity and during spontaneous activity. On the other hand, the same IMFs would be obtained if the EMD process were applied to the whole recording. Indeed, we should highlight that the IMF for spontaneous activity and after whisker stimulation are centered around the same frequency because we employ a mask sifting algorithm. Mask sifting is a variant of Empirical Mode Decomposition that reduces mode mixing by introducing a sinusoidal masking signal during the sifting process [3, 4]. The mask defines a reference frequency scale that guides the adaptive separation of oscillatory components, rather than imposing a strict bandpass filter. This method is particularly well-suited for intermittent oscillations, as observed in our experiments, where classical EMD approaches are known to encounter difficulties. Indeed, in such conditions, standard EMD relies entirely on local extrema, making the decomposition highly sensitive to amplitude fluctuations and temporal irregularity. Selecting the first mask at 30 Hz (which is the one we pass to the algorithm), the effective lower bound of the first oscillatory content is approximately $0.7 \times 30$ Hz, i.e. about 21 Hz [3]. In this sense, the first intrinsic mode function predominantly contains activity in the range starting from $\sim$21 Hz and above, while components with lower characteristic frequencies are progressively shifted to subsequent IMFs or to the residual. In standard implementations, mask frequencies are typically arranged in a dyadic sequence, i.e. scaling by a factor of 2 (termed *mask-step-factor*) the previous one (the masks would then be 30 Hz, 15 Hz, 7.5 Hz, etc.). This corresponds to successive effective lower bounds of approximately 21 Hz, 10.5 Hz, 5.25 Hz, 2.62 Hz and so on, producing a hierarchical decomposition from higher to lower frequency content and improving scale separation compared to classical EMD. In this regard, it is important to emphasize that the selection of mask frequencies does not impose the presence of oscillatory components at those frequencies. The masks act as reference scales that guide the decomposition and improve scale separation, without introducing artificial spectral content. Consequently, an intrinsic mode function exhibits high power at a given frequency only if that component is genuinely present in the signal. This can be observed, for example, in the IMFs extracted from the thalamus signals: although they are peaked in the same frequency range as those of the cortex, their power is substantially lower for IMF2, which is the relevant aspect for our analysis.

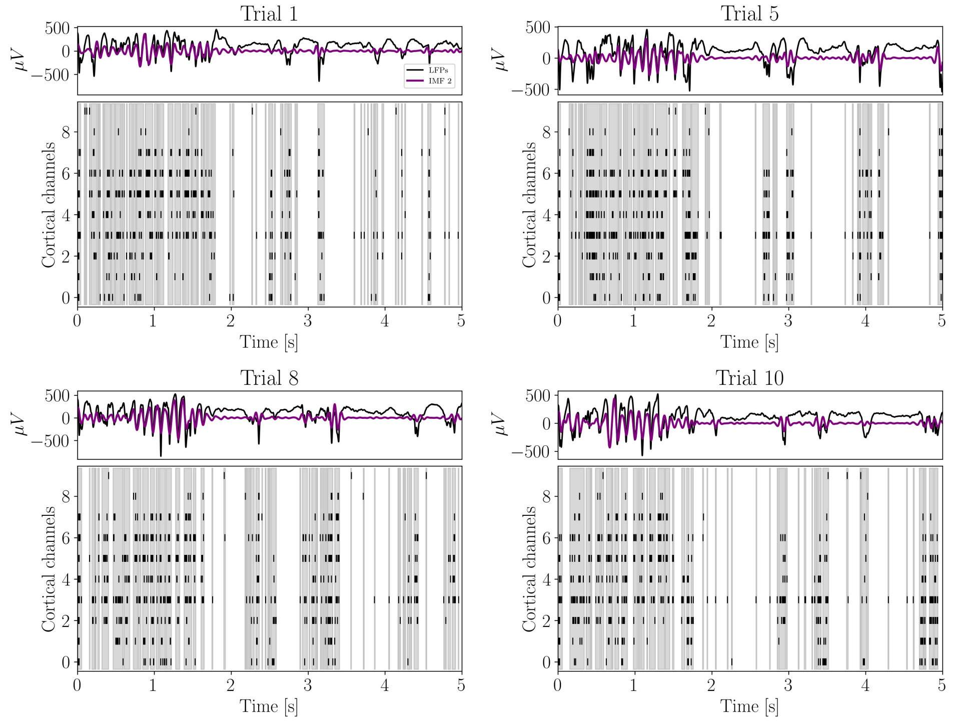


FIG. S3: **Relationship between the fast oscillation and avalanches**. 4 examples of activity after stimulation of the whisker are shown (top panels in each subplot: LFPs signals and IMF 2 signals. bottom panels: raster plot obtained from MUAs and detected avalanches (empty bins indicate the end of an avalanche).

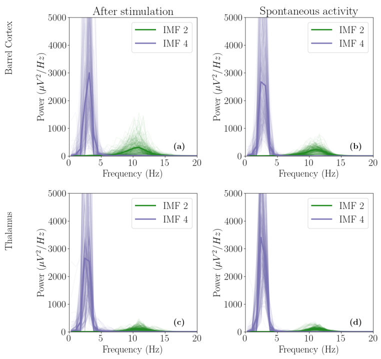


(1)

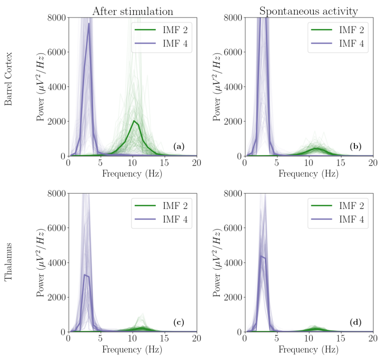


(2)

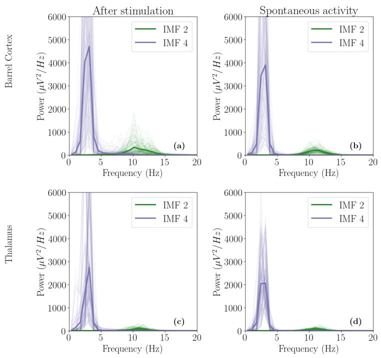


(3)

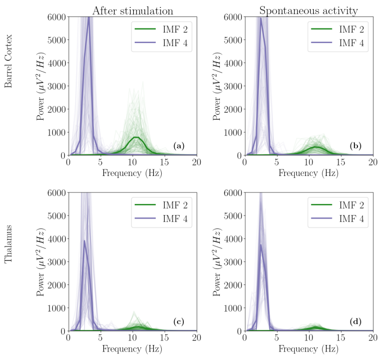


(4)

FIG. S4: Marginal Hilbert spectrum of intrinsic Mode Functions 2 and 4 in each of the four rats analyzed (**1**, **2**, **3**, **4**). **(1)** (a) Marginal Hilbert spectrum after the whisker stimulation in the rat barrel cortex. (b) Marginal Hilbert spectrum during spontaneous activity in the rat barrel cortex. (c) Marginal Hilbert spectrum after the whisker stimulation in the rat thalamus. (d) Marginal Hilbert spectrum during spontaneous activity in the rat thalamus. Same in **2**, **3**, **4**, for three additional rats.

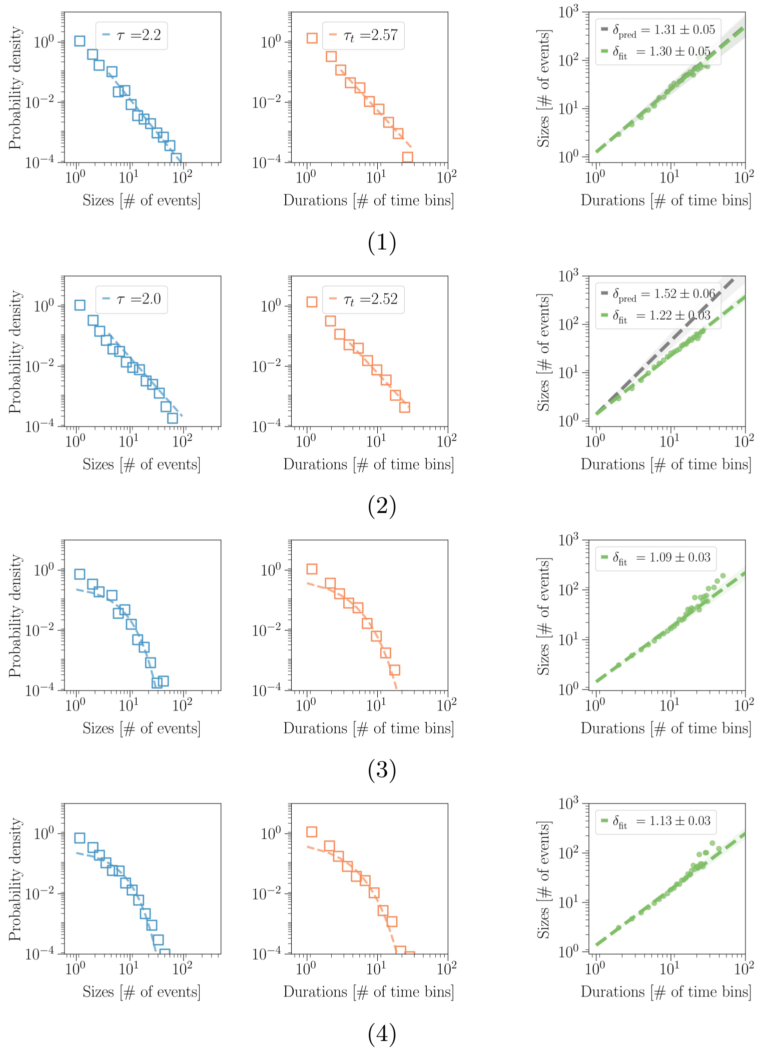


FIG. S5: Avalanches distribution in the barrel cortex during spontaneous activity (1) and after stimulation of the whisker (2) Avalanches distribution in the thalamus during spontaneous activity (3) and after stimulation of the whisker (4) (Rat 1)

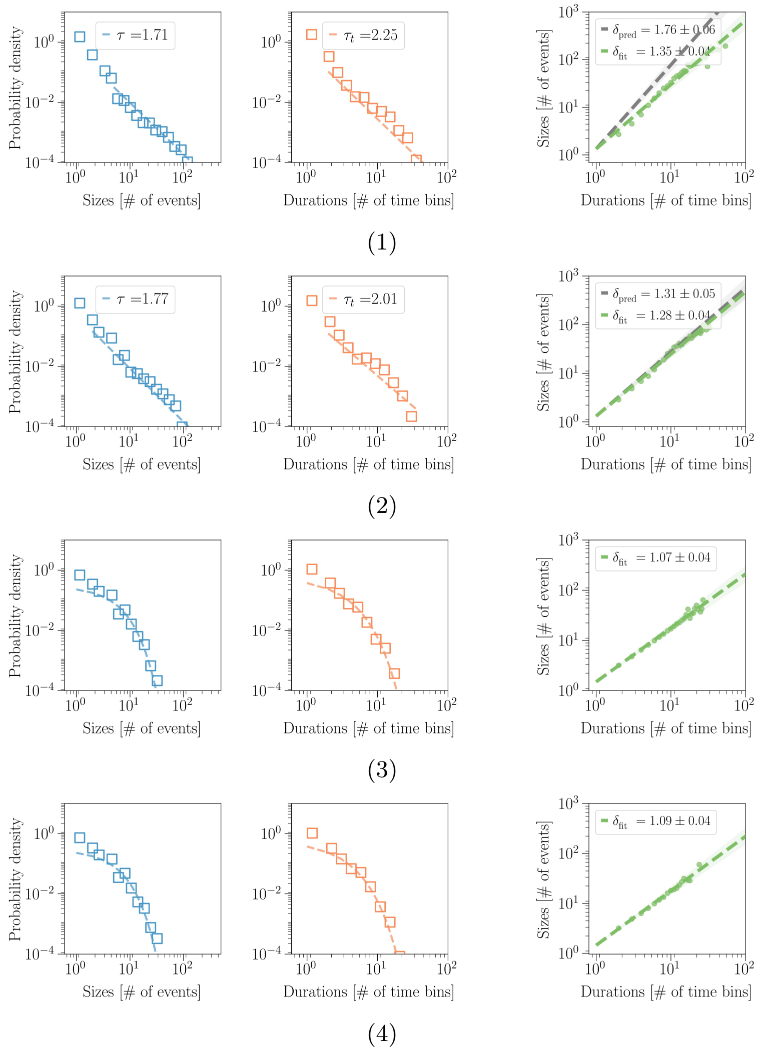


FIG. S6: Avalanche distributions for an additional rat (rat 2), same analysis as in Figure S5

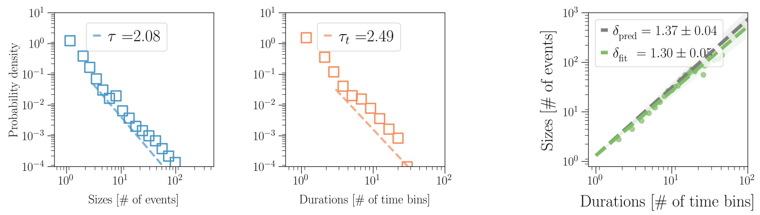


(1)

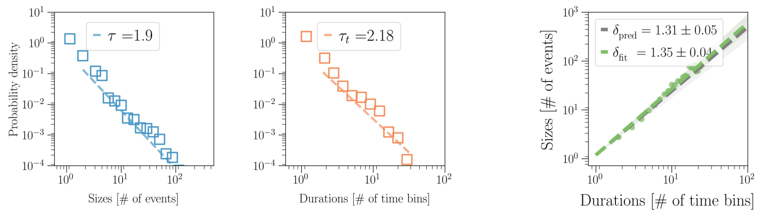


(2)

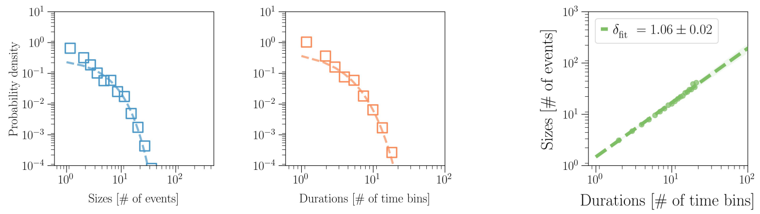


(3)

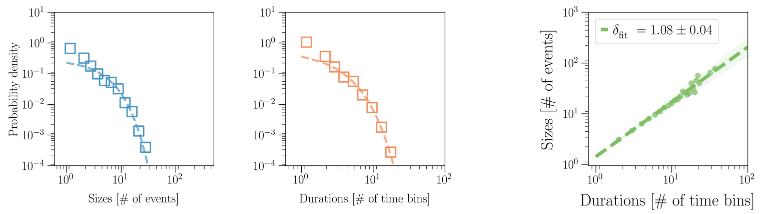


(4)

FIG. S7: Avalanche distributions for an additional rat (rat 3), same analysis as in Figure S5

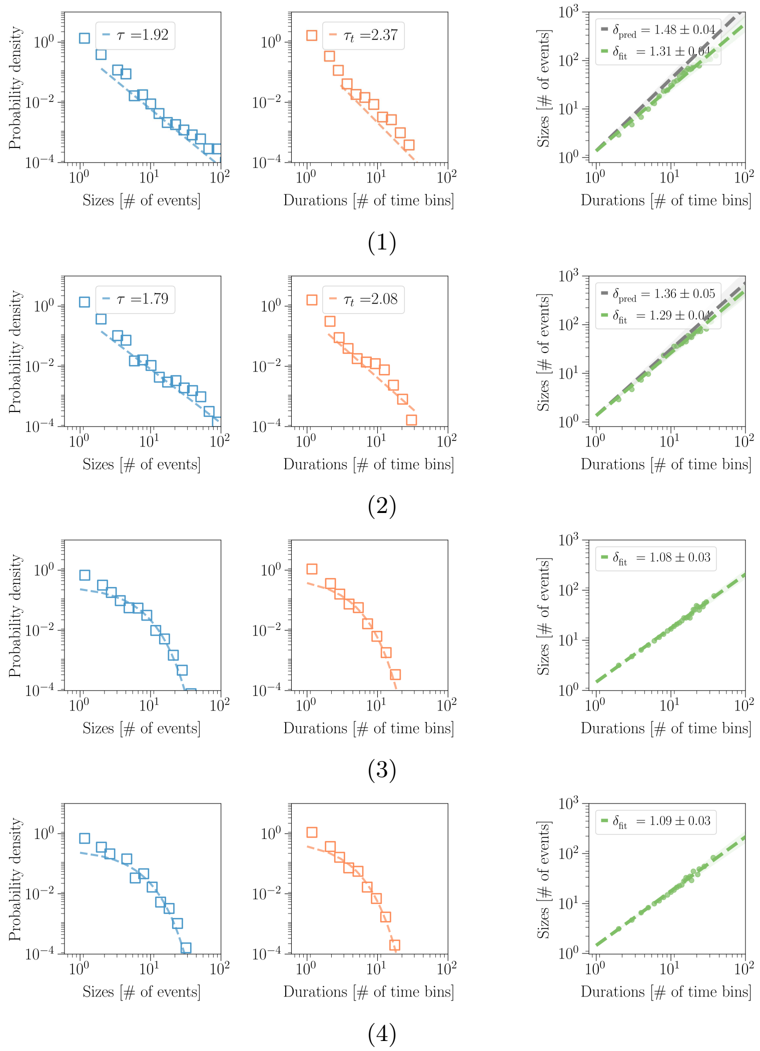


FIG. S8: Avalanche distributions for an additional rat (rat 4), same analysis as in Figure S5

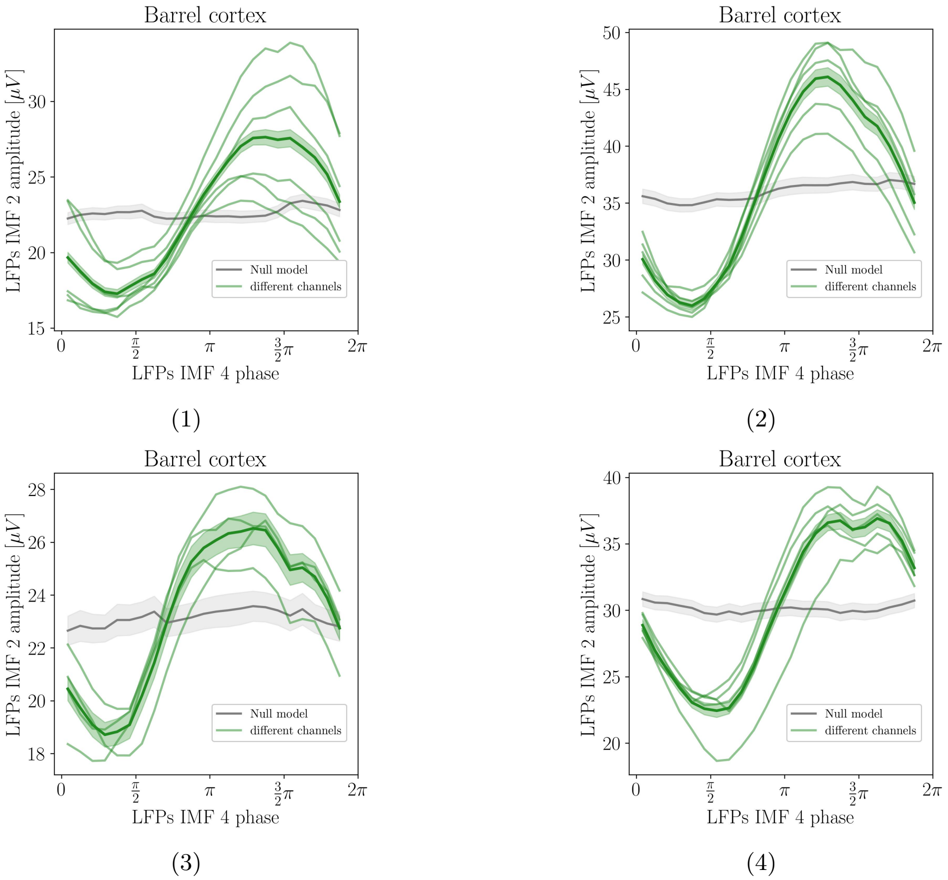


FIG. S9: Phase amplitude coupling (PAC) between the phase of intrinsic mode function 4 and the amplitude of intrinsic mode function 2, in four different rats ((1),(2),(3),(4)). The dark green line indicates the average PAC across channels, while lighter green lines indicate single-channel results.

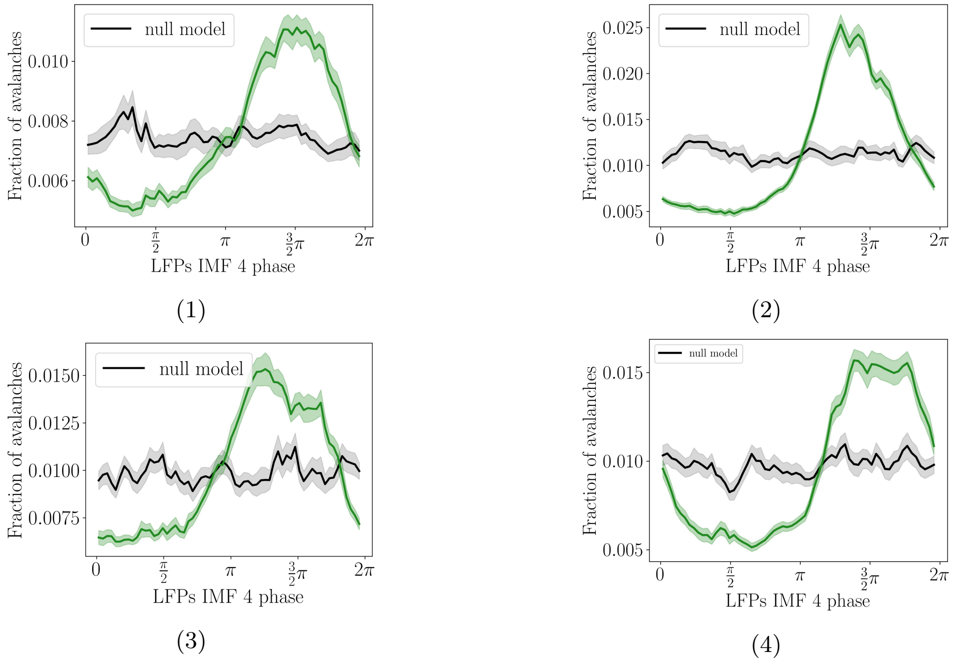


FIG. S10: Coupling between the phase of Intrinsic mode function 4 and the fraction of avalanches, in four different rats ((1),(2),(3),(4)). The null model (in black) is computed by randomly shuffling the trials relative to the IMF 4 signals, i.e., disrupting the temporal correlation between the IMF 4 phase and avalanches.

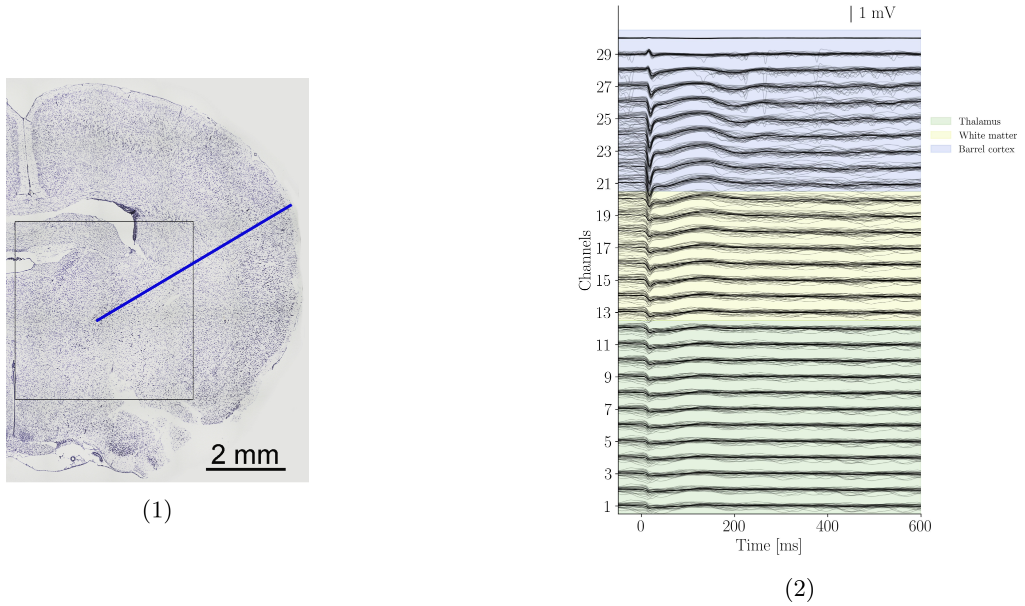


FIG. S11: **Histological verification of probe insertion and LFPs profile** (1) Brain right hemisphere slice stained with cresyl violet. The blue trace indicates the probe insertion path. The black box is centered on the thalamus. Scale bar: 2 mm. (2) LFPs profile in response to whisker stimulation (at time 0 s), across the cortex, white matter, and thalamus as measured by the neural probe [5, 6]. As usual, the placement of the probe is evaluated before starting the recordings by stimulating different whiskers following the somatotopic map existing between the mystacial pad and the S1 cortical architecture, to identify the whisker that gives the strongest evoked response and thus to identify the corresponding barrel.

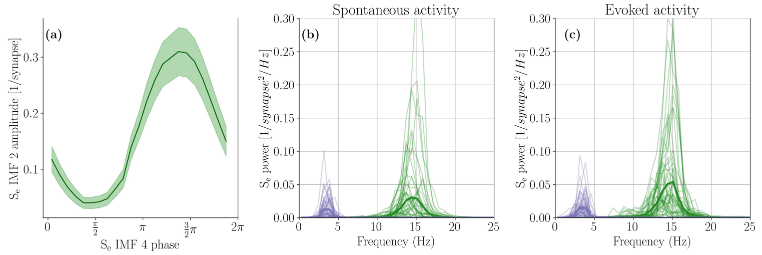


(1)

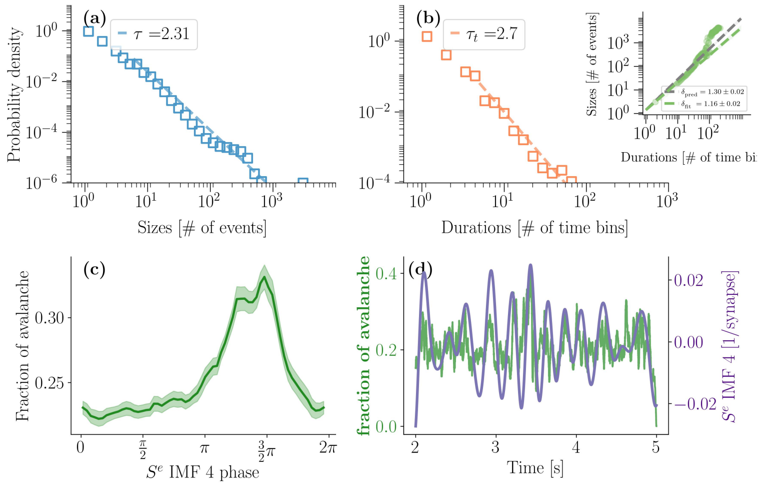


(2)

FIG. S12: **Results for the model (Rat 1)** The dynamics of the model that receives as input the thalamic firing rate from Rat 1 is shown here. (1) Oscillatory activity of the model during spontaneous activity (b) and evoked activity (c). Phase amplitude coupling between the two modes is shown in (a). (2) Avalanche sizes (a) and durations distributions (b) in the model. In (c, d), the coupling of the fraction of avalanches with the phase of IMF 4 is shown.

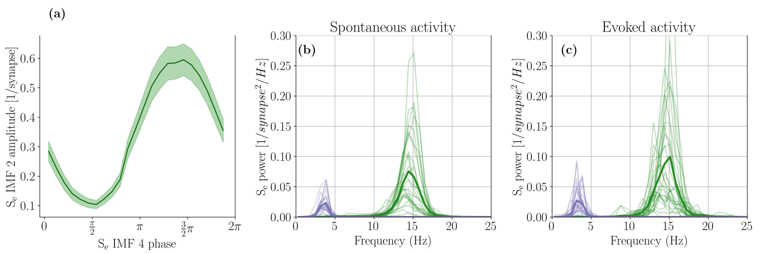


(1)

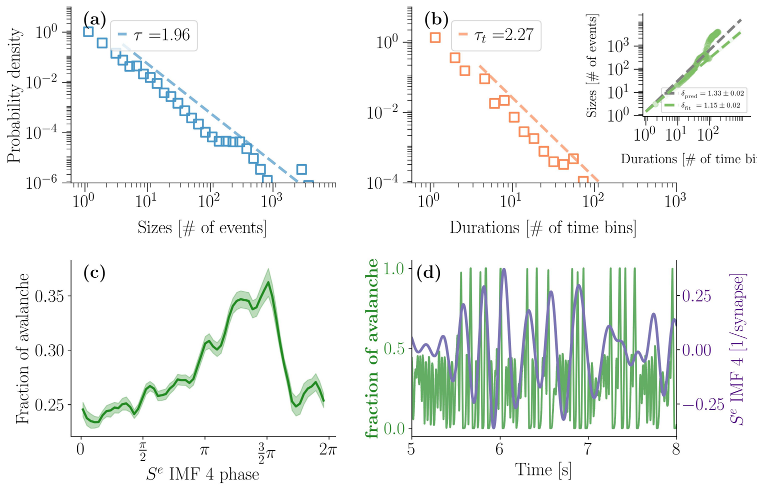


(2)

FIG. S13: **Results for the model (Rat 2)** Same as in S12, for rat 2.

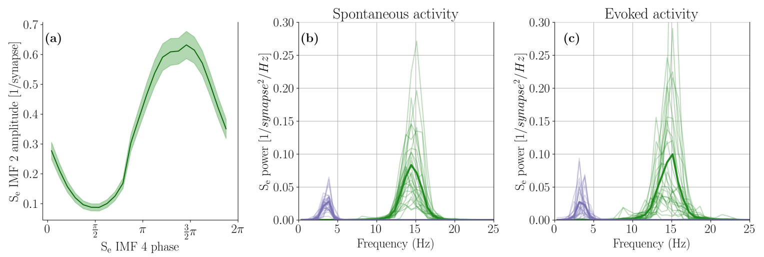


(1)

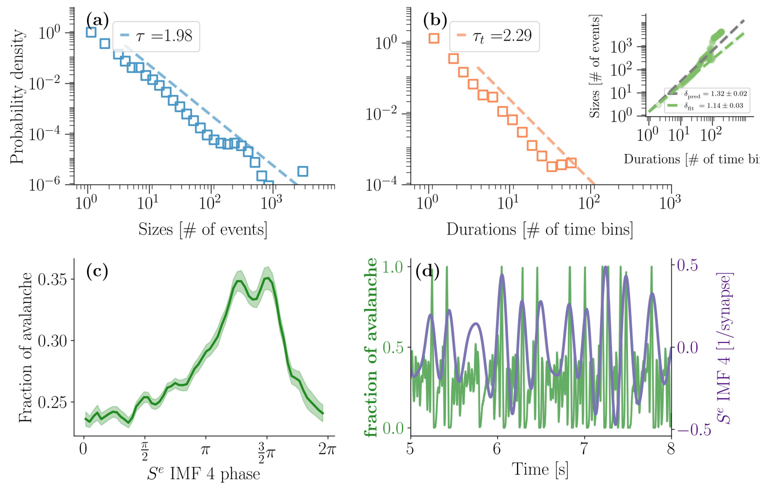


(2)

FIG. S14: **Results for the model (Rat 3)** Same as in S12, for rat 3.

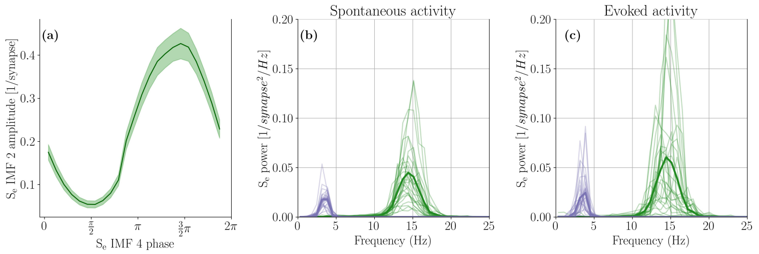


(1)

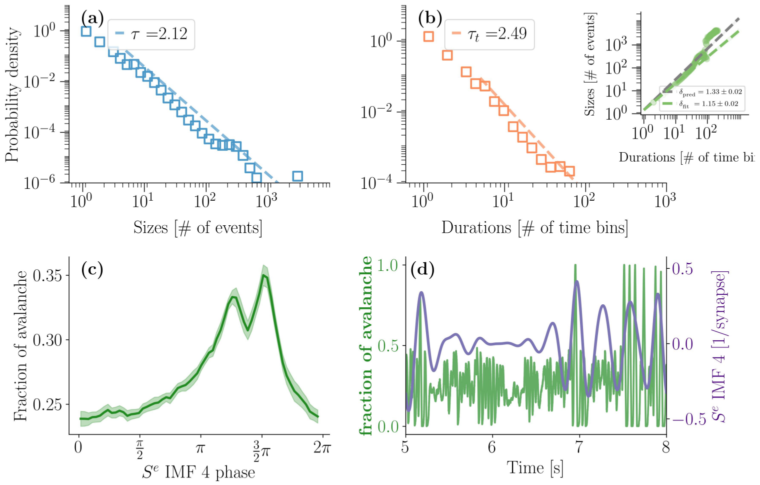


(2)

FIG. S15: **Results for the model (Rat 4)** Same as in S12, for rat 4.

[1] Mike X Cohen. *Analyzing neural time series data: theory and practice.* MIT press, 2014.

[2] Vincent Douchamps, Matteo Di Volo, Alessandro Torcini, Demian Battaglia, and Romain Goutagny. Gamma oscillatory complexity conveys behavioral information in hippocampal networks. *Nature Communications*, 15(1):1849, 2024.

[3] Andrew J. Quinn, Vitor Lopes-dos Santos, David Dupret, Anna C. Nobre, and Mark W. Woolrich. Emd: Empirical mode decomposition and hilbert-huang spectral analyses in python. *Journal of Open Source Software*, 6(59):2977, 2021.

[4] Ryan Deering and James F Kaiser. The use of a masking signal to improve empirical mode decomposition. In *Proceedings.(ICASSP'05). IEEE International Conference on Acoustics, Speech, and Signal Processing, 2005.*, volume 4, pages iv–485. IEEE, 2005.

[5] Simona Temereanca and Daniel J Simons. Local field potentials and the encoding of whisker deflections by population firing synchrony in thalamic barreloids. *Journal of neurophysiology*, 89(4):2137–2145, 2003.

[6] Hanno S Meyer, Robert Egger, Jason M Guest, Rita Foerster, Stefan Reissl, and Marcel Oberlaender. Cellular organization of cortical barrel columns is whisker-specific. *Proceedings of the national academy of sciences*, 110(47):19113–19118, 2013.
